# Passage of Zika virus in *Rag1*-deficient mice selects for unique envelope glycosylation motif mutants that show enhanced replication

**DOI:** 10.1101/2024.09.06.611742

**Authors:** Eri Nakayama, Bing Tang, Romal Stewart, Abigail L. Cox, Kexin Yan, Cameron R. Bishop, Troy Dumenil, Wilson Nguyen, Andrii Slonchak, Julian Sng, Alexander A. Khromykh, Viviana P. Lutzky, Daniel J. Rawle, Andreas Suhrbier

**Author notes:** Co first authors (Eri Nakayama, Bing Tang). Co-last authors (Daniel J. Rawle, Andreas Suhrbier).

## Abstract

N-linked glycosylation of flavivirus envelope proteins is widely viewed as a requirement for optimal folding, processing and/or transit of envelope proteins, and the assembling virons, through the endoplasmic reticulum (ER) and Golgi. Herein we show that serial passage of ZIKV_Natal_ in *Rag1*^-/-^ mice generated two unique envelope glycan-deficient mutants, ZIKV-V153D and ZIKV-N154D, that, surprisingly, produced titers ∼1 to 2.6 logs <u>higher</u> than the glycosylated parental ZIKV_Natal_ in Vero E6 cells and human brain organoids. RNA-Seq of infected organoids suggested that this increased replication fitness was associated with upregulation of the unfolded protein response (UPR). Cell death, cellular viral RNA and viral protein levels were not significantly affected, arguing that these glycan mutants enjoyed faster ER/Golgi folding, processing, assembly, transit, and virion egress, assisted by an upregulated UPR. Thus, ZIKV envelope N-linked glycosylation is not essential for promoting envelope folding, assembly and transit through the ER/Golgi, as aspartic acid (D) substitutions in the glycosylation motif achieve this with significantly greater efficiency. V153D and N154D mutants have not been employed in flavivirus envelope glycosylation studies. Instead, mutants such as N154A have been used, which may impart unfavorable properties that have a greater impact than the loss of the glycan. ZIKV-V153D and -N154D may avoid this by preserving the surface negative charge provided by the glycan moiety in the parental ZIKV_Natal_. In *Ifnar*^-/-^ mice ZIKV-V153D and -N154D showed faster viremia onsets, but reduced viremic periods, than the parental ZIKV_Natal_, consistent with the contention that these glycans have evolved to delay neutralizing antibody activity.

**IMPORTANCE:** Studies seeking to understand the role(s) of N-linked glycosylation of flavivirus envelope proteins often introduce amino acid substitutions that disrupt the glycosylation motif, which in ZIKV has the sequence ^153^VNDT^156^. Unfortunately, such substitutions, for instance N154A, may themselves impart unfavorable properties on envelope that have a greater impact than the loss of the glycan moiety. Herein we describe two unique glycosylation motif mutants, ZIKV-V153D and -N154D that were positively selected during passage of ZIKV_Natal_ in *Rag1*^-/-^ mice. These N^154^ glycan-deficient viruses produced viral titers up to ∼400 fold higher than the parental ZIKV_Natal_ in Vero cells and in human brain organoids. Such glycans are thus clearly not a requirement for optimal folding and trafficking of ZIKV envelope through the endoplasmic reticulum/Golgi. These results provide new insights into the molecular mechanisms underpinning viral fitness *in vitro* and *in vivo*, and also have implications for virus-like-particle vaccine design and production.

## INTRODUCTION

Zika virus (ZIKV) is a positive sense, single-stranded RNA arbovirus that belongs to the *Flavivirus* genus (renamed *Orthoflavivirus* in 2023) in the family Flaviviridae, and is primarily transmitted to humans by *Aedes aegypti* and *Aedes albopictus* mosquitoes. ZIKV emerged as an important pathogen in 2007, with outbreaks in the Pacific Islands and subsequent spread across the Americas in 2015-2016 (1). The World Health Organization (WHO) regards ZIKV as a priority disease for research and development after declaring a public health emergency of international concern in 2016. ZIKV infection of pregnant women can lead to congenital Zika syndrome (CZS), which comprises a spectrum of primarily neurological disorders in neonates (2, 3). CZS can be associated with severe functional impairments that have profound adverse impacts on child development and quality of life (4). After ZIKV is introduced by the bite of an infected mosquito, viral dissemination and viremia is primarily driven by infection of subsets of monocytes and macrophages (5). ZIKV can then cross the placenta and enter the developing fetal brain, perhaps via transcytosis across the endothelial cells comprising the blood brain barrier (6, 7). Once in the brain, ZIKV can infect a number of brain cell types, including neural progenitor cells, a critical cell population in the developing embryonic brain. Infection of these cells results in growth perturbations, inhibition of differentiation and apoptosis, all key processes in driving the brain damage associated with CZS (8, 9).

The envelope glycoprotein of ZIKV, and other flaviviruses, is responsible for viral entry into host cells and is a key determinant of viral pathogenesis (10-13), as well as being the target of protective neutralizing antibodies (14). The envelope proteins of nearly all flaviviruses have N-linked glycosylation(s), which for ZIKV is a single glycosylation site located at amino acid N^154^. The ZIKV envelope glycan moiety is adjacent to the fusion loop in the mature virion, and has been proposed to reduce access of neutralizing antibodies targeting the fusion loop region (15-17). However, glycan moieties on viral envelope proteins are reported to have a range of activities (18, 19), with a large body of literature arguing that loss of N-linked glycans on flaviviral envelope proteins reduces *in vitro* replication, often leading to reduced virulence in animal models (Supplementary Table 1). Other studies show no change or mixed effects (Supplementary Table 1). The endoplasmic reticulum (ER) is arguably the critical hub for flavivirus replication (20); and a key mechanism proposed to underpin the aforementioned observations is that N-linked glycosylation of the flavivirus envelope protein supports proper folding, processing, stability, maturation and/or transport of envelope, and the assembling virion, during the journey through the ER and Golgi, and the eventual egress into the extracellular milieu (19-24). In contrast, a number of publications report increased replication and/or virulence associated with the loss of N-linked glycosylation of flaviviral envelope proteins (25-29) or find no, or mixed, effects (13, 16, 30-34) (Supplementary Table 1). The disparity between these studies may arise, at least in part, from the variety of strategies used to disrupt the N-glycosylation motif (Supplementary Table 1), which for ZIKV usually has the amino acid sequence ^153^VNTD^156^ (35), although inclusion of V^153^ in this motif has only been proposed (36-38). A number of flavivirus studies have used N154A substitutions to prevent N-link glycosylation at this site (22, 35, 39, 40), whereas others have used different substitutions or deletions in the glycosylation motif (13, 26, 41, 42) (Supplementary Table 1). A problem of interpretation emerges as either the loss of the glycan moiety and/or the change in the envelope amino acid sequence may be responsible for the effects on virus replication. For instance, the N154A substitution has been associate with reduced solubility of ZIKV envelope protein and an ensuing rapid degradation by the host ER-associated degradation system (22). The N154A substitution also increased the toxicity of ZIKV envelope protein expressed in neuronal cells *in vitro* (43). Thus for the latter, it is unclear whether the loss of the glycan is responsible, and/or whether the loss of the glycan is secondary to introduction of the small hydrophobic amino acid, alanine, that itself imparts unfavorable properties on the envelope protein.

ZIKV replication in wild-type mice is inefficient, with mouse model work thus often involving use of IFNα/β receptor deficient (*Ifnar^-/-^*) mice (9, 44-46). We previously reported infection of *Ifnar^-/-^* mice with an African lineage virus isolate, maintained at the National Institute of Infectious Diseases (Tokyo, Japan), ZIKV-MR766-NIID (7). This virus isolate does not have an N-linked glycosylation in the envelope protein due to a D156I substitution in the glycosylation motif, VND<u>I</u>. However, within 4 days post infection (dpi), the virus recovered from the serum of *Ifnar^-/-^* mice showed restoration of the glycosylation motif (VNDT) (7). *Ifnar^-/-^* mice are fully capable of generating neutralizing antibody responses against ZIKV (46, 47). IgM can also neutralize ZIKV (48, 49) and antigen-specific IgM can appear as early as 1-3 dpi (50-52). This selection for the N-linked envelope glycan may thus have arisen to avoid neutralizing antibody responses (15-17). However, selection may also have occurred because the glycan is required for optimal folding, processing and/or traffic of envelope through the ER. Also possible is that the glycan moiety is selected because it promotes infection of host cells via lectin attachment factors (53).

To generate more insights into the role of the ZIKV N-linked envelope glycan, and perhaps inform the aforementioned issues, we sought to determine what changes might be selected in mice in the absence of antibody responses. *Rag1*^-/-^ mice were chosen as they are unable to mount adaptive immune responses, but can nevertheless support ZIKV replication (54), despite intact type I IFN responses. Murine type I IFN responses effectively blunt ZIKV replication (55, 56); however, antibodies are needed to clear the viremia (57). We chose to use ZIKV_Natal_ (58) as this Asian lineage isolate is unequivocally associated with a human case of microcephaly and has only a brief passage history in C6/36 cells (59), whereas ZIKV-MR766 viruses have been extensively passaged in young mice and are usually more virulent (7, 60). Viral titers reached by ZIKV_Natal_ infections are generally lower than for ZIKV-MR766 (7, 46), with deep mutational scanning data also available for ZIKV_Natal_ (45). Herein, ZIKV_Natal_ was passaged five times in *Rag1*^-/-^ mice in three independent passage series. Remarkably, each passage series selected a virus with mutations in the envelope protein glycosylation motif. Two of these, ZIKV-V153D and ZIKV-N154D, generated titers ∼1-2.6 logs higher than the glycosylated parental ZIKV in Vero E6 cells and in human cortical brain organoids. These results argue against a critical role for glycans for folding, processing, assembly and/or transit of envelope through the ER, as this was clearly achieved with much greater efficiency by appropriate amino acid substitutions. In *Ifnar^-/-^* mice, viremia peaked earlier, but the viremia period was shorter, for the two glycan mutants, when compared with the parental ZIKV_Natal_. These observations favor the view that these glycans have evolved to delay generation of, and targeting by, neutralizing antibody responses (61, 62) that are generated within a few days post infection.

## MATERIALS AND METHODS

### Ethics statement and regulatory compliance

All mouse work was conducted in accordance with the “Australian code for the care and use of animals for scientific purposes” as defined by the National Health and Medical Research Council of Australia. Mouse work was approved by the QIMR Berghofer Medical Research Institute Animal Ethics Committee (P2195, A1604-611M and P3746, A2108-612). Mice were euthanized using CO_2_. Breeding and use of GM mice was approved under a Notifiable Low Risk Dealing (NLRD) Identifier: NLRD_Suhrbier_Oct2020: NLRD 1.1(a). All work was approved by the QIMR Berghofer Medical Research Institute Safety Committee (P2195 and P3746). Details of mouse agistment conditions have been described in detail previously (63).

### Cell lines and viral titrations

Vero cells (ATCC#: CCL-81) and C6/36 cells (ATCC# CRL-1660) cell were maintained in RPMI 1640 (Thermo Fisher Scientific, Scoresby, VIC, Australia) supplemented with endotoxin free 10% heat-inactivated fetal bovine serum (FBS; Sigma-Aldrich, Castle Hill, NSW, Australia) at 37°C and 5% CO_2_. Cells were checked for mycoplasma using MycoAlert Mycoplasma Detection Kit (Lonza, Basel, Switzerland).

ZIKV titers were determined by focus forming unit (ffu) assays (45), or by CCID_50_ assays using ten-fold serial dilution titrations, in duplicate in 96 well plates as described (7). For the latter, virus titers were determined by the method of Spearman and Karber (64).

### Generation of ZIKV_Natal_

The complete genome sequence for ZIKV_Natal_ was obtained from fetal brain of a human case of microcephaly (59) (GenBank accession number KU527068). Infectious virus was reconstructed using reverse genetics to produce a primary ZIKV_Natal_ isolate, with no mouse passage history (58). ZIKV_Natal_ stocks were produced in C6/36 cells as described (58).

### ZIKV_Natal_ passaging in *Rag1*^-/-^ mice

Adult *Rag1^-/-^* mice (B6.129S7-^Rag1tm1Mom^/J, JAX) bred in-house were infected with ∼10^6^ ffu of ZIKV_Natal_. Mice were weighed 3 times a week, and when weight loss reached at least 15%, mice were euthanised and blood was taken via heart puncture into MiniCollect serum separation tubes (Greiner Bio-One GmbH, Kremsmunster, Austria). Serum was used for virus titrations by CCID_50_ assays, and 100 µl of serum was passaged into a new *Rag1*^-/-^ mice by intravenous inoculation. Mouse brains were also collected and tissue titers determined by CCID_50_ assays as described (7).

### Nanopore sequencing of ZIKV from *Rag1*^-/-^ mouse serum, end of passage 5

ZIKV RNA was isolated from mouse serum using Nucelospin RNA virus kit (Macherey-Nagel, Düren, Germany), and cDNA was synthesized using ProtoScript II Reverse Transcriptase (New England Biolabs) as per manufacturer’s instructions. Barcoding PCR was performed with ZIKV-specific primers containing the Nanopore universal tail sequences of 5′-TTTCTGTTGGTGCTGATATTGC-3′ for the forward primer and 5′-ACTTGCCTGTCGCTCTATCTTC-3′ for the reverse primer. Primers were designed to amplify approximately 1 kb overlapping amplicons spanning the entire ZIKV genome. PCR using pooled primers (Supplementary Table 2; odd and even-numbered primers pooled separately in two reactions to avoid primer interference at the overlapping sequences) was performed using Q5 High-Fidelity 2X Master Mix (New England Biolabs) as per manufacturer’s instructions with the following thermocycling conditions; 98°C 30 s, 5 cycles of 98°C 10 s, 55°C 30 s, 72°C 1 min, then 25 cycles of 98°C 10 s, 65°C 30 s, 72°C 1 min, and final extension of 72°C 2 min. Amplicons were gel-purified using QIAquick Gel Extraction Kit (QIAGEN). The two amplicon pools for each sample were further pooled and the second round of barcoding PCR was performed using LongAmp Taq 2X Master Mix (New England Biolabs) and Oxford Nanopore barcoding primer sets BC07, BC08, BC09 and BC11, each containing a unique barcode. DNA repair and end-prep using NEBNext FFPE DNA Repair Mix and NEBNext Ultra™ II End Repair/dA-Tailing Module (New England Biolabs) was performed as per manufacturer’s instructions. Adapter ligation and clean-up were performed using NEBNext Quick Ligation Module (New England Biolabs) as per manufacturer’s instructions. Sequencing runs were conducted using the MinION Flongle flow cell using MinKNOW software (Oxford Nanopore Technologies, UK) as described (65). Guppy basecaller (V4.0.11; https://nanoporetech.com/) was used to convert .fast5 files to fastq files. Alignments were undertaken using minimap2 (v2.16) and ZIKV_Natal_ (GenBank: KU527068) as the reference genome. The Integrative Genomics Viewer v2.8.0 (Broad Institute, USA) was used to visualize sequence data.

### Virus stock generation for ZIKV-V153D and ZIKV-N154D

Virus stocks for ZIKV-V153D and ZIKV-N154D were generated by 2 passages in C6/36 cells. Initially, C6/36 cells were inoculated with serum from viremic *Rag1*^-/-^ mice (taken at euthanasia after passage 5), and were cultured for 5/6 days. Supernatants were then transferred to fresh C6/36 cells and supernatants harvested on 6 dpi, stored at -80°C in aliquots, and then used as virus stocks for subsequent experiments. The glycosylation motif regions from all stocks were sequenced by capillary sequencing.

### Capillary sequencing

To confirm ZIKV RNA sequences in particular regions after virus stock propagation in C6/36 cells or after infection of *Ifnar*^-/-^ mice, viral RNA was harvested using Nucelospin RNA virus kit (Macherey-Nagel, Düren, Germany), and cDNA was synthesized using ProtoScript II Reverse Transcriptase (New England Biolabs) as per manufacturer’s instructions. PCR was performed using primers in Supplementary Table 2 that capture the region of interest, or primers to amplify the envelope N-linked glycosylation site region; forward 5’-CTACCTTGACAAGCAATCAG-3’ and reverse 5’-CCAACCAGTGCTTGTTATTC-3’. PCR products were gel-purified using Monarch DNA Gel Extraction Kit (New England Biolabs) as per manufacturer’s instructions and sequences determined by BigDye Terminator v3.1 Sanger sequencing using either the forward or reverse primer. Capillary sequencing was then undertaken using SeqStudio 8 Flex Genetic Analyzer (Applied Biosystems).

### PNGase F treatment and western blot

Vero E6 cells were infected with ZIKV_Natal_, ZIKV-V153D or ZIKV-N154D stock viruses. Cell monolayers were washed with ice-cold PBS before addition of 200 µl trypsin (Sigma) to detach the cells. Cells were collected and centrifuged at 500 g at 4°C for 5 min, and washed with ice-cold PBS and centrifuged again at 500 g, 4°C for 5min. Cells were resuspended in lysis buffer (50 mM Tris HCl, 150 mM NaCl, 1mM EDTA, 1% v/v Triton X-100, pH 7.4) with protease inhibitor cocktail (cOmplete™ Protease Inhibitor Cocktail, Merck Cat# 11697498001) and incubated on ice for 20 min. The lysate was centrifuged at 13,000 g for 20 min and the supernatant was harvested and aliquots stored at -80°C. Twenty µg of protein was treated with PNGase F as per manufacturer instructions (New England Biolabs). Water was used as the mock treatment. The reactions were stopped by adding SDS-PAGE sample buffer (125 mM Tris–HCl, pH 6.8; 4% [v/v] SDS; 20% [v/v] glycerol; 0.004% [w/v] bromphenol blue) and boiling for 10 min. Samples were analyzed using 10% SDS-PAGE gel and Western blotting using 4G2 (1:100) as the primary antibody incubated at 4°C overnight. HRP goat anti-mouse was use as the secondary antibody (1:2000), and chemiluminescence substrate (SuperSignal™ West Pico PLUS Chemiluminescent Substrate) was used as per manufacturers’ instructions. Blots were imaged using Biorad ChemiDoc Touch Imaging System.

### Growth kinetics in C6/36 or Vero E6 cells

C6/36 and Vero E6 cells were infected in triplicate with the indicated ZIKV stocks at a multiplicity of infection (MOI) of 0.01. C6/36 cells were seeded at 2×10^5^ cells in 24 well plate and were incubated at 28°C. Vero E6 were seeded at 10^6^ in 6 well plates and were incubated at 37°C. After one hour the supernatants were removed and the cells washed in PBS three times before adding fresh culture media (RPMI 1640 supplemented with 10% fetal calf serum). Supernatants were collected at the indicated times and virus titers determined by CCID_50_ assays.

### Human brain organoids; generation, infection and monitoring

The human-induced pluripotent cells (hiPSCs) utilized in this study were reprogrammed from adult dermal fibroblasts (HDFa, Gibco, C0135C) employing the CytoTune-iPS 2.0 Sendai Reprogramming Kit (Invitrogen, A16518; (66)). These cells were maintained on human recombinant vitronectin-coated plates (Thermo Fisher Scientific) in StemFlex medium (Thermo Fisher Scientific), according to the manufacturer’s guidelines.

Human cortical brain organoids were generated as outlined (65, 67, 68). hiPSCs were dissociated using StemPro Accutase (Thermo Fisher Scientific) to produce a single-cell suspension. The cells were then plated at a density of 5,000 cells per well in an ultra-low-binding 96-well plate (Corning) containing StemFlex medium supplemented with 10 μM ROCK inhibitor Y-27632 (STEMCELL Technologies, Vancouver, Canada). From days 1 to 5, the media was replaced daily with StemFlex medium supplemented with 2 μM Dorsomorphine (Abcam) and 10 μM SB-431542 (Stemcell Technologies). On day 5, the medium was replaced with a Neuro-induction medium comprising DMEM/F12 (Thermo Fisher Scientific), 1% N2 Supplement (Thermo Fisher Scientific), 10 μg/mL heparin (STEMCELL Technologies), 1% penicillin/streptomycin (Thermo Fisher Scientific), 1% Non-essential Amino Acids (Thermo Fisher Scientific), 1% Glutamax (Thermo Fisher Scientific), and 10 ng/mL FGF2 (Stemcell Technologies). On day 7, the organoids were embedded in Matrigel (Corning), transferred to an ultra-low-binding 24-well plate (Corning) (one organoid per well), and cultured in Neuro-induction medium for an additional three days. On day 10, the organoids were transitioned to a differentiation medium comprising Neurobasal medium, 1% N2, 2% B27 supplements (Thermo Fisher Scientific), 0.5% Penicillin/Streptomycin, 1% Glutamax, 1% Non-essential Amino Acids, 50 μM 2-mercaptoethanol (Merck), 2.5 μg/mL Insulin (Merck), 1% Knockout Serum Replacement (Thermo Fisher Scientific), 10 ng/mL FGF2, and 1 μM CHIR99021 (Stemcell Technologies). The organoids were cultured for an additional five days with media changes performed every other day.

Fifteen-day-old cortical organoids, measuring ∼1 to 1.3 mm in diameter, were infected in quadruplicate with 5×10^5^ CCID_50_ of either ZIKV_Natal_, ZIKV-V153D, or ZIKV-N154D stock viruses for 4 hours at 37°C in a 5% CO_2_ environment. The organoids were then washed twice with media and cultured in neuro-differentiation media at 37°C, 5% CO_2_ (67, 69). Supernatants were collected at the indicated time points, and viral titres determined by CCID_50_ assays as described (7).

Organoids were imaged at various intervals using an EVOS FL microscope (Advanced Microscopy Group), and their 2D image circumference was measured using Image J v1.53 by outlining the organoid periphery (70). At 4 dpi, organoids (n=4 per group) were harvested and formalin-fixed for histology and immunohistochemistry.

### Immunohistochemistry of human brain organoids

The immunostaining was conducted using the pan-flavivirus 4G4 (anti-NS1) and 4G2 (anti-envelope) monoclonal antibodies as described (68, 71) using 3 organoids per virus, with organoids harvested at 4 dpi. Briefly, organoids were embedded in agarose prior to standard paraffin embedding. Sections were stained with 4G4 or 4G2 and HRP-conjugated Perkin Elmer goat anti-mouse secondary antibody, with signal developed using Vector Nova Red. Sections were lightly counterstained with hematoxylin. Slides were scanned using Aperio AT Turbo (Aperio, Vista, CA USA) and analyzed by and Aperio Positive Pixel Count Algorithm (Leica Biosystems) that provided weak positive, positive and strong positive brown staining pixel counts (Nova Red) divided by total count of blue staining pixels (hematoxylin counterstain).

### RNA-Seq and bioinformatics

Total RNA was extracted from organoids using TRIzol (Invitrogen). RNA concentration and quality were measured using TapeStation D1kTapeScreen assay (Agilent). cDNA libraries were generated using Illumina TruSeq Stranded mRNA library prep kit, and were sequenced using an Illumina Nextseq 2000 Sequencing System to produce 75 nt paired-end reads. Sequence reads were aligned to the ZIKA virus genome (KU527068.1) with bowtie2 v2.2.9 (72). Primary proper read alignments to the viral genome were counted using Samtools v1.10 (73). Sequence reads were aligned to the GRCh38 human reference genome obtained from Gencode, using STAR aligner v2.7.10 (74). Aligned read counts were obtained for annotated genes using RSEM v1.3.1 (75), and differential expression was estimated using DESeq2 v1.32.0 with R statistical software v4.2.0 ((76) and R Core Team). Significantly differentially expressed genes (DEGs) were analysed using Ingenuity Pathway Analysis v107193442 (QIAGEN). Gene set enrichment analyses (GSEA) were performed using GSEA v4.3.2 with gene sets from the Molecular Signatures database v2023 (77) and the All gene list ranked by fold change.

For single nucleotide variant (SNV) detection, viral read alignments were pre-processed to re-align indels, mark duplicates, and recalibrate basecall quality scores, using lofreq v2.1.5 (78) and the Genome Analysis Toolkit v4.2.4.1 (79). Variants were called using Lofreq with an intra-host allele frequency cutoff of greater than 0.15 and a p-value cutoff of less than 0.05. Intra-host allele frequencies were averaged across replicates and read-depth per SNV was summed across replicates.

### Infection of *Ifnar*^-/-^ mice

Female *Ifnar*^-/-^ mice (12-13 weeks old) (originally provided by Prof P. Hertzog, Monash University, Melbourne, VIC, Australia (80)) on a C57BL/6J background (81) were bred in-house and were infected intraperitoneally with 10^3.2^ CCID_50_ of ZIKV_Natal_, ZIKV-V153D or ZIKV-N154D virus stocks that had been generated in C6/36 cells. Mice were weighed daily and bled via the tail vein for viremia determinations using CCID_50_ assays.

### Statistics

The t-test (unequal variance) was used (in Microsoft Excel 2016, Data Analysis tool) when the difference in variances was <4, skewness was > -2 and kurtosis was <2. Otherwise, the non-parametric Kolmogorov-Smirnov Exact test was used in GraphPad Prism 8.2.1.

## RESULTS

### Passage of ZIKV_Natal_ in *Rag1*^-/-^ mice

ZIKV_Natal_ was used to infect *Rag1*^-/-^ mice, which have no mature B or T lymphocytes, with viremia and weight loss subsequently monitored. The animals were euthanized when the ethically defined end point of >15% body weight loss was reached. Serum from euthanized mice was then passaged into new *Rag1*^-/-^ mice. This process was repeated for five passages in *Rag1*^-/-^ mice, with passaging conducted in three replicate series comprising three independent parallel mouse-to-mouse transfer series (5 mice per series) (Fig. 1A). The cumulative amount of time that *Rag1*^-/-^ mice were infected with ZIKV_Natal_ ranged from 180 days (Replicate #3 series) to 400 days (Replicate #2 series) (Fig. 1B). The time to the weight-loss endpoint (>15%) and euthanasia tended to decrease after 1-2 passages and stabilized at approximately 30-40 days for the remaining 3 passages in all three replicates (Fig. 1C).

**Figure 1.**
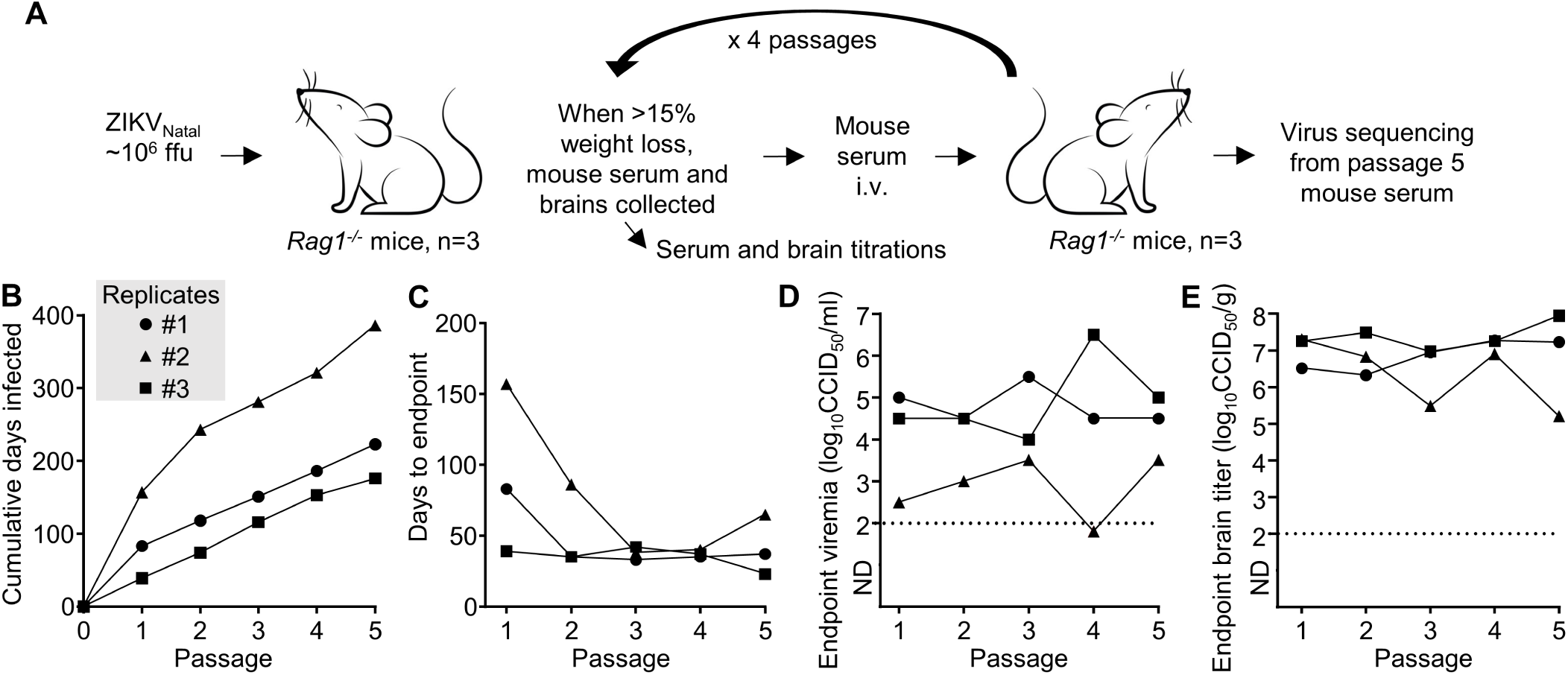
Passage of ZIKV_Natal_ in *Rag1*^-/-^ mice. (A) Experimental design schematic. Three *Rag1*^-/-^ mice were initially infected with ZIKV_Natal_, when weight loss reached >15%, mice were euthanized and serum passaged to new *Rag1*^-/-^ mice; repeated 5 times in 3 replicate passage series (n=15 mice in total). (B) For each of the 3 replicate passage series, the cumulative number of days that the virus was present/replicated in *Rag1*^-/-^ mice. (C) For each of the 3 replicate passage series, the number of days between virus inoculation and when weight loss reached >15%, requiring the animal to be euthanized. (D) For each of the 3 replicate passage series, viremia for each mouse at euthanasia. Limit of detection ∼2 log_10_CCID_50_/ml. (E) For each of the 3 replicate passage series, brain virus titers for each mouse at euthanasia.

The serum titers in each *Rag1*^-/-^ mouse over the initial 2-6 dpi for each of the Replicate series were relatively low for the first 1-2 passages, thereafter (passages 3-5) they increased and remained at ∼2-3 log_10_CCID_50_/g (Supplementary Fig. 1). The serum titers at euthanasia did not show a clear trend with passage number (Fig. 1D), with arboviral viremias in *Rag1^-/-^*mice tending towards a steady state level after 1-2 weeks (82). However, one mouse had an undetectable viremia at passage 4 (limit of detection was ∼2 log_10_CCID_50_/ml); nevertheless, serum transfer successfully established a detectable viremia at passage 5 (Fig. 1D).

Brain ZIKV_Natal_ titers at euthanasia were consistently between 5 - 8 log_10_CCID_50_/g (Fig. 1E), indicating that there was no significant increase in brain infection due to serial passage. Brain infection and associated weight loss in ZIKV-infected *Rag1*^-/-^ mice have been described previously (83).

### ZIKV passage in *Rag1*^-/-^ mice selects for envelope N154-linked glycosylation motif mutants

After 5 passages in *Rag1*^-/-^ mice, the virus in the serum from euthanized mice (end of passage 5) for each of the 3 replicate series was sequenced using Nanopore sequencing. Sequences were aligned to the ZIKV_Natal_ reference genome and nucleotide changes ≥ 15% were manually curated using Integrated Genomics Viewer (IGV) (Supplementary Table 3). Across the three replicates, there were 42 total nucleotide mutations resulting in approximately 50% synonymous and 50% non-synonymous amino acid mutations (Supplementary Table 3). The resulting amino acid changes are shown in Fig. 2A. In all three independent replicates, the envelope N-linked glycosylation motif was disrupted, with this involving 3 different patterns of nucleotide and amino acid changes (Fig. 2B). The ZIKV_Natal_ N-linked envelope glycosylation motif comprises the amino acids VNDT at positions 153-156. The dominant sequence changes in this motif were Replicate #1 54% VDDT and 41% DNDT, Replicate #2 VDDT, and Replicate #3 INDI (Fig. 2B).

**Figure 2.**
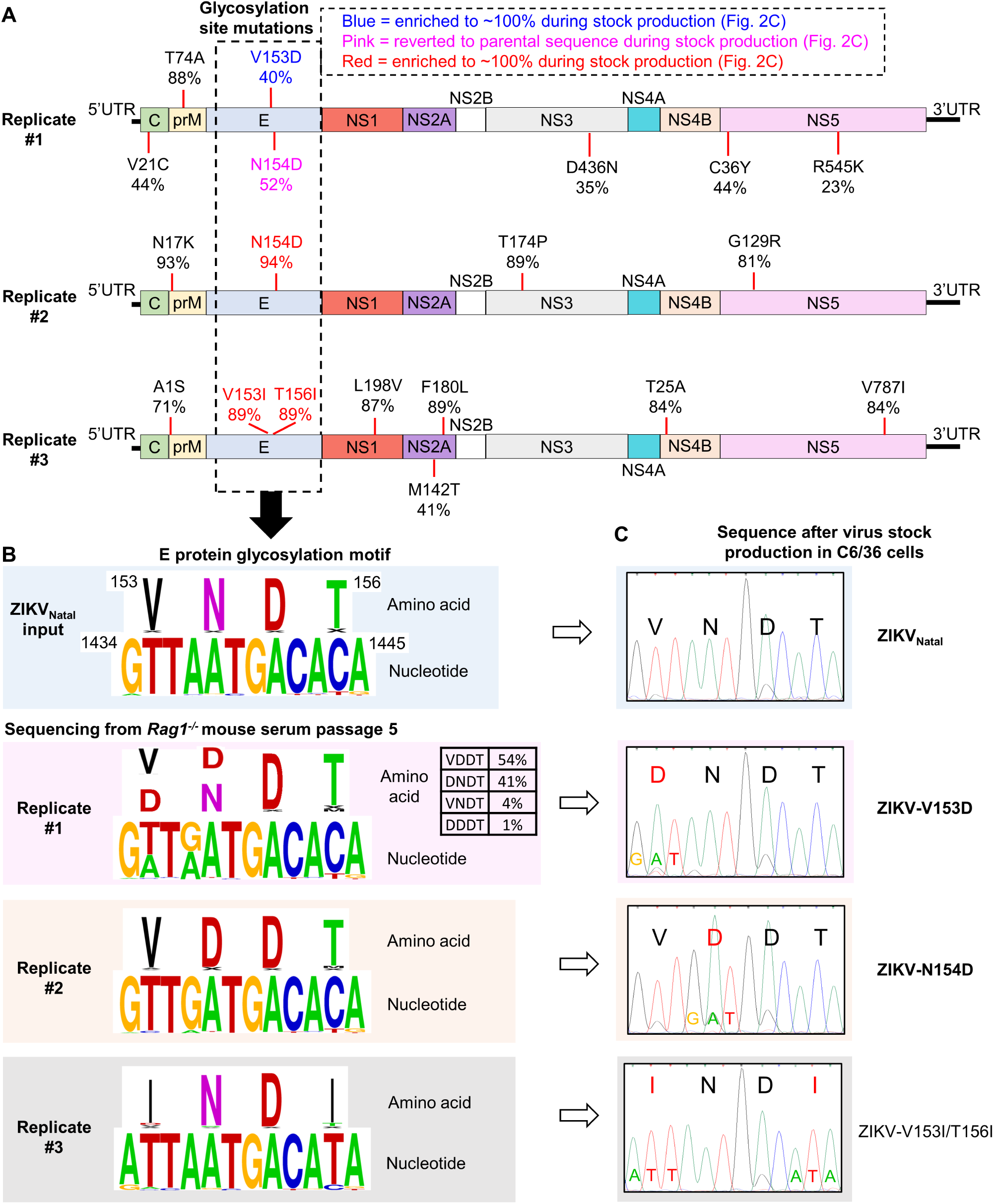
Sequences of ZIKV_Natal_ after passage in *Rag1*^-/-^ mice. (A) Amino acid changes in the ZIKV_Natal_ sequences of viruses in serum of *Rag1*^-/-^ mice after passage 5 for each of the three replicate passage series. Sequences were obtained by Nanopore sequencing. Reference genome KU527068.1. (B) Amino acid and nucleotide sequences of the envelope N-linked glycosylation motif of the parental ZIKV_Natal_ and the viruses described in A. Amino acid and nucleotide numbering from KU527068.1. (C) Envelope N-linked glycosylation motif sequences after generation of virus stocks by 2 passages in C6/36 cells. Sequences were obtained by capillary sequencing of PCR products (Supplementary Fig. 2A, B).

Viruses from *Rag1*^-/-^ mouse serum (end of passage 5) were passaged twice in C6/36 cells to produce virus stocks, with capillary sequencing of the glycosylation site illustrating that disruptions in the N-linked glycosylation motif were retained in the virus stocks (Fig. 2C; Supplementary Fig. 2A, B). Although Replicate #1 presented a mixed population in *Rag1*^-/-^ mouse serum (Fig. 2A, B), the DNDT sequence emerged as dominant after passage in C6/36 cells (Fig. 2C, ZIKV-V153D), with T74A and D436N also retained (Supplementary Fig. 2C).

In summary, during serial passage of ZIKV_Natal_ in *Rag1*^-/-^ mice, disruptions of the envelope N-linked glycosylation motif were identified in all 3 independent replicates. Each disruption was distinct and glycosylation motif disruptions were retained after generation of viral stocks in C6/36 cells (Fig. 2C; Supplementary Fig. 2A, B).

### PNGase treatment to assess N-linked glycosylation of ZIKV envelope proteins

To confirm that the aforementioned mutations disrupt the N-linked glycosylation of envelope, C6/36-derived viral stocks were used to infect Vero E6 cells, and at 4 dpi, cells were harvested and cell lysates treated with PNGase F and analysed by Western blotting using the 4G2 anti-E monoclonal antibody. The reduction in molecular weight was evident for envelope from ZIKV_Natal_ (Fig. 3A, white arrow). No reductions in the molecular weight of envelope from ZIKV-V153D, ZIKV-N154D or ZIKV-V153I/T156I were evident after PNGase F treatment (Fig. 3A, white dashed line), arguing that the envelope proteins from these viruses do not have an N-linked glycan.

**Figure 3.**
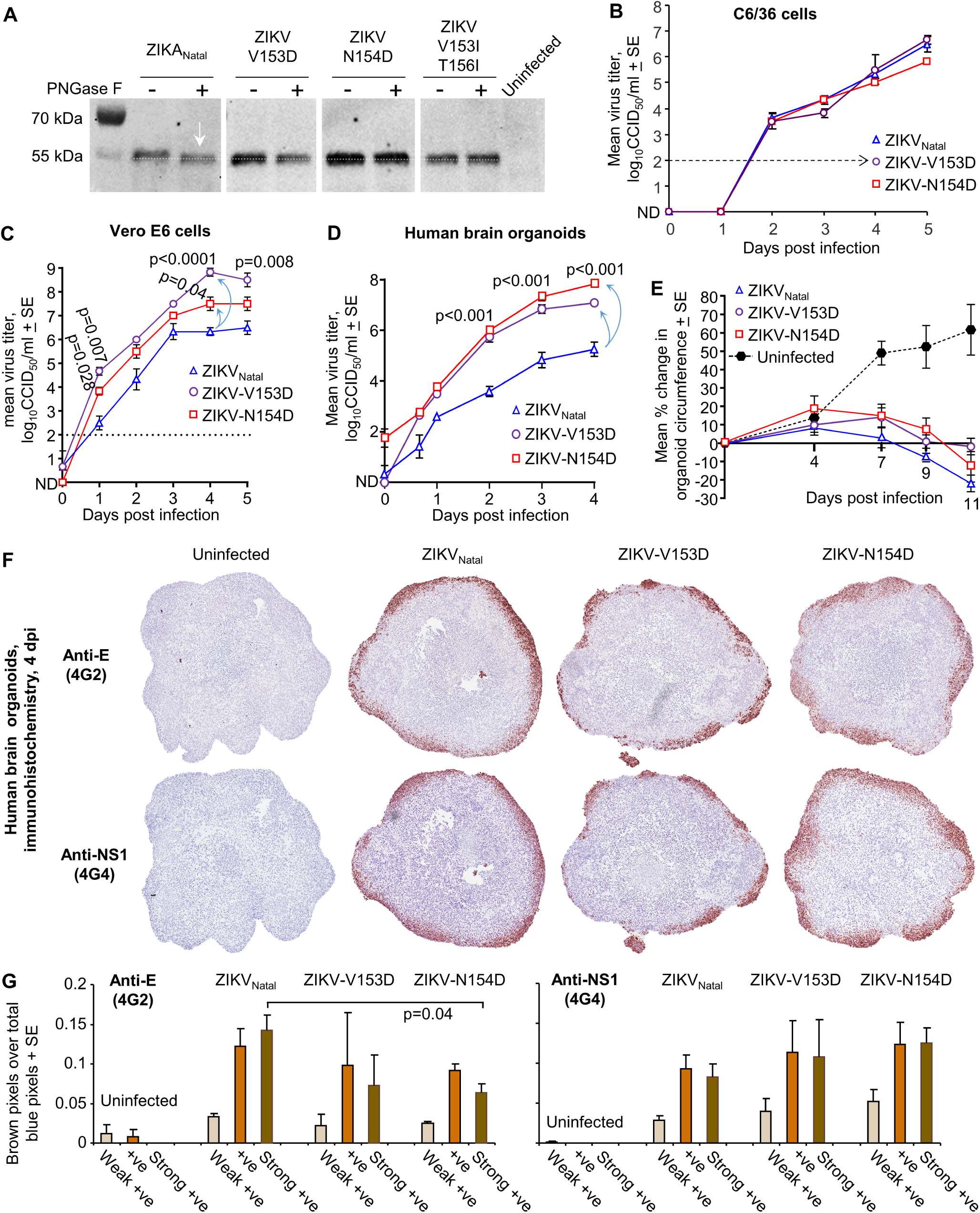
Glycosylation and replication of glycosylation motif mutants. The following experiments were conducted using C6/36-derived virus stocks. (A) The indicated viruses were used to infect Vero E6 cells and cell lysates (4 dpi) were treated with PNGase F and analyzed by Western blotting using the 4G2 anti-E monoclonal antibody. The reduction in molecular weight is evident for ZIKV_Natal_ (white arrow). (B) Growth kinetics of the indicated (C6/36 cell-derived) virus stocks in C6/36 cells. Limit of detection ∼2 log_10_CCID_50_/ml. (C) Growth kinetics of the indicated (C6/36 cell-derived) virus stocks in Vero E6 cells. At 4 dpi the mean titer for ZIKV-V153D was ∼2.5 logs higher, and the mean titer for ZIKV-N154D was ∼1 log higher, than ZIKV_Natal_ (light blue arrows). Statistics by t test, all vs. ZIKV_Natal_. (D) Growth kinetics of the indicated (C6/36 cell-derived) virus stocks in 15 day old human cortical brain organoids (mean titers were derived from 4 replicates for each virus). At 4 dpi the mean titers for ZIKV-V153D were ∼1.8 logs higher, and for ZIKV-N154D the mean titers were ∼2.6 logs higher, than ZIKV_Natal_ (light blue arrows). Statistics by t test vs. ZIKV_Natal_. (E) Mean percentage change in organoid circumference at 11 dpi vs. 0 dpi for each organoid (n=4 organoids per group). (F) IHC of brain organoids at 4 dpi using pan-flavivirus anti-NS1 (4G4) and anti-E (4G2) monoclonal antibodies. The diameter of the uninfected 15 day old organoids was ∼1-1.3 mm. (G) Aperio pixel count analysis of antibody staining (weak positive – light brown), positive (brown), strong positive (dark brown), divided by total blue staining (hematoxylin counterstain). Means shown for n=3. Statistics by t test.

### ZIKV N-linked glycosylation mutants show increased replication in Vero E6 cells and human brain organoids

Replication kinetics of C6/36-derived virus stocks were assessed in C6/36 cells. Growth of ZIKV-V153D, ZIKV-N154D and ZIKV_Natal_ were very similar in C6/36 cells (Fig. 3B). ZIKV-V153I/T156I replicated very poorly in C6/36 cells (Supplementary Fig. 3), and was thus not used in further experiments. In Vero E6 cells, ZIKV-V153D and ZIKV-N154D both replicated to significantly higher titers (∼2.5 and 1 logs, respectively) when compared to the parental ZIKV_Natal_ (Fig. 3C).

Human brain organoids have been used extensively to study ZIKV infection of neuronal cells (65, 84). Early (15 day old) human cortical brain organoids were infected with C6/36-derived stocks of ZIKV_Natal_, ZIKV-V153D and ZIKV-N154D, and viral titers in culture supernatants determined over time. Consistent with data from Vero E6 cells (Fig. 3C), ZIKV-V153D and ZIKV-N154D both replicated to significantly higher titers (∼1.8 and 2.6 logs, respectively, at 4 dpi) compared to ZIKV_Natal_ in the human brain organoids (Fig. 3D, blue arrows). Curiously, reductions in organoid circumferences, driven by ZIKV cytopathic effects (CPE) (65), were not significantly higher for ZIKV-V153D or ZIKV-N154D, when compared with ZIKV_Natal_ (Fig. 3E). Furthermore, immunohistochemistry (IHC) of organoids at 4 dpi did not indicate significantly increased levels of envelope (4G2) or NS1 (4G4) protein staining for ZIKV-V153D or ZIKV-N154D infected organoids, when compared with ZIKV_Natal_ infected organoids (Fig. 3F, G), despite the logs higher levels of virus replication at 4 dpi (Fig. 3D). Pixel count analysis of strong brown 4G2 staining was actually slightly, but significantly, lower for ZIKV-N154D infected organoids (Fig. 3G, p=0.04).

In summary, ZIKV-V153D and ZIKV-N154D showed significantly increased replication (up to ∼2.6 logs) in Vero E6 cells and human brain organoids when compared with ZIKV_Natal._ IHC and size analyses of infected human brain organoids indicated that the log increases in replication were not associated with infection of more cells, increased viral envelope or NS1 protein expression, or substantially increased levels of CPE.

### RNA-Seq suggests more robust induction of the unfolded protein response by ZIKV-N154D

To gain insights into the processes that might underpin the increased replication of the glycan mutants (Fig. 3D), ZIKV-N154D infected human cortical organoids were compared with ZIKV_Natal_ infected organoids at 4 dpi by RNA-Seq. ZIKV-N154D was chosen as the N154D mutation is unequivocally associated with loss of N-linked glycosylation, as the glycan is attached to the side chain of N154.

The number of viral RNA reads was not significantly different for ZIKV-N154D vs. ZIKV_Natal_ (Fig. 4A), despite the log increases in the levels of infectious virus in the supernatants of these organoids (Fig. 3D). Taken together, these data suggest that the steady state viral RNA levels in the infected cells are largely unchanged, but that mature virion egress was substantially accelerated. Analysis of the viral sequences (Supplementary Fig. 4) illustrated retention of the N154D substitution for ZIKV-N154D and no changes in the glycosylation motif for ZIKV_Natal_ (Fig. 4B).

**Figure 4.**
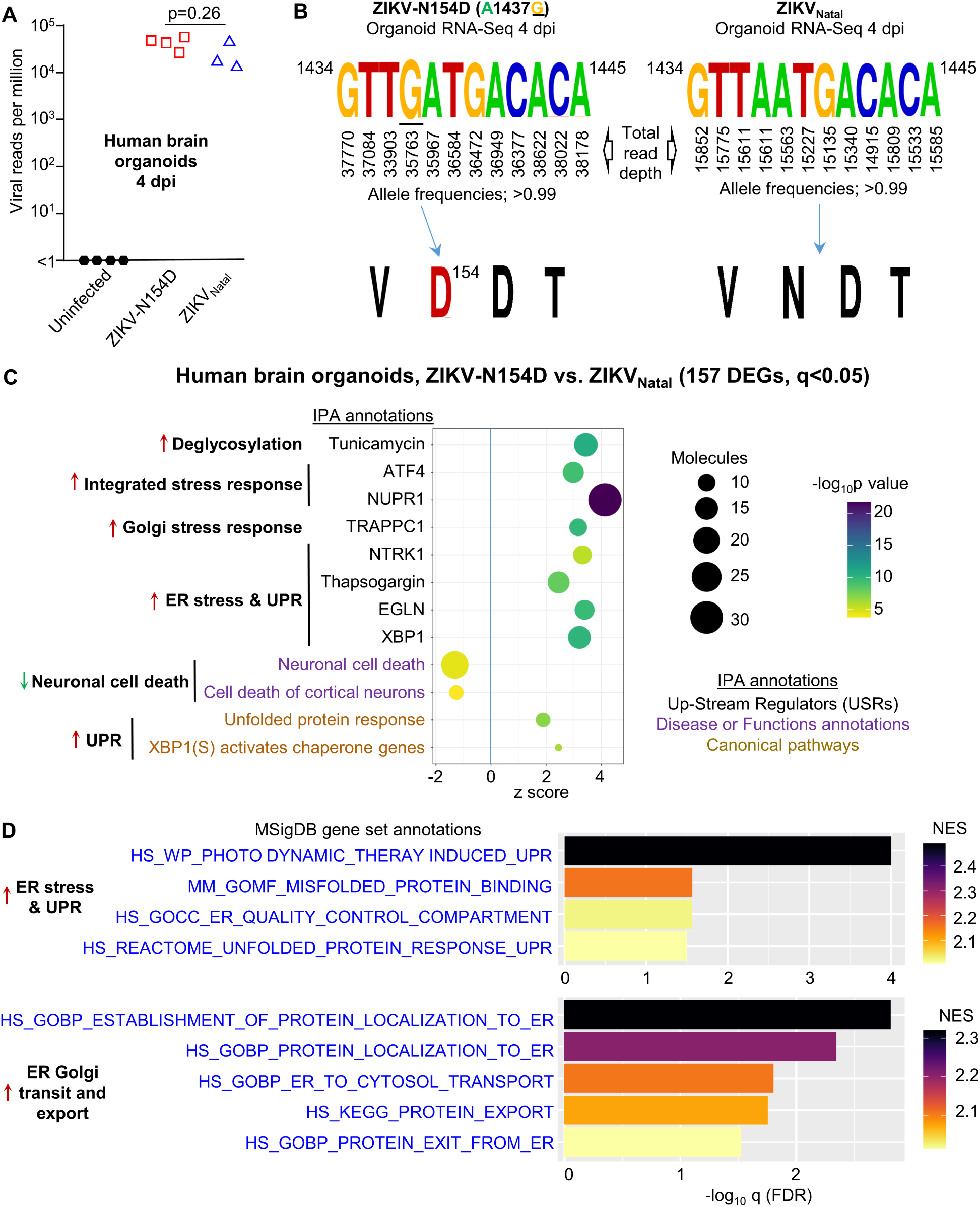
Brain organoid RNA-Seq. (A) Viral read counts per million at 4 dpi for organoids infected with the indicated virus. Statistic by t test (not significant). (B) Left - retention of N154D substitution for the ZIKV-N154D after 2 passages in C6/36 cells and 4 days growth in organoids. Right – no nucleotide changes were identified after 4 days growth of ZIKV_Natal_ in organoids. The read depth at each position is derived from the combined data from all organoids infected with the indicated virus. (C) Ingenuity Pathway Analyses (IPA) of DEGs identified by RNA-Seq data comparing ZIKV-N154D with ZIKV_Natal_ infected organoids at 4 dpi. Selected annotations are shown and are grouped into themes (bold text), with arrows indicating up-regulation (red) or down-regulation (green). (D) GSEA analysis of ranked All gene list using MSigBD gene sets, selected annotations are shown as for C. Full list of genes and bioinformatic analyses are provided in Supplementary Table 4.

Analyses of the host cell transcriptome provided 157 differentially expressed genes (DEGs, q<0.05) for ZIKV-N154D vs. ZIKV_Natal_ (Supplementary Fig. 5, Supplementary Table 4), with these DEGs then used in Ingenuity Pathway Analyses (IPA). Tunicamycin was identified as significant upstream regulator (USR) (Fig. 4C), perhaps expected as this is a drug that inhibits N-linked glycosylation. Annotations associated with the Golgi stress response, TRAPPC1 (85) and the integrated stress response, NUPR1 (86) and ATF4 (87), were also identified. Several IPA annotations indicated an increase in ER stress and the unfolded protein response (UPR). These included XBP1 (88-90) and Thapsogargin (91), with NTRK1 (92) and EGLN (93) also associated with the UPR (Fig. 4C). IPA Diseases and Functions annotations also emerged indicating lower levels of neuronal cell death in ZIKV-N154D infected organoids when compared with ZIKV_Natal_ infected organoids (Fig. 4C, Supplementary Table 4). This is consistent with a marginal trend toward less CPE for ZIKV-N154D vs. ZIKV_Natal_ as indicated by organoid size, although this did not reach significance (Fig. 3E).

GSEAs were undertaken using the ranked All gene list (Supplementary Table 4) and gene sets from the MSigDB. A series of annotation associated with ER stress/UPR were again identified (Fig. 4D) consistent with the IPA data. A series of MSigDB annotations also suggested increased protein traffic through the ER and Golgi, and out of the cell (Fig. 4D, ER Golgi transit and export). Such activities would be consistent with a successful UPR, which leads to upregulation of secretory pathway chaperones and foldases that reduce the nascent protein load in the ER (94) by increasing protein export (95-97). A successful UPR can also mitigate against cell death (94).

In summary, the RNA-Seq data argues that the glycan mutant, ZIKV-N154D, induces a more robust and successful UPR than the glycosylate parent ZIKV_Natal_, which results in increased transit of the viral envelope protein, and the assembling virion, through the ER and Golgi and out of the cell. Such activity would be consistent (96, 97) with the increased viral titers seen in Fig. 3D, without significant increases in cytoplasmic viral or viral protein levels (Fig. 3F, G).

### ZIKV-V153D and ZIKV-N154D replicate faster in *Ifnar^-/-^* mice

To determine whether the increased replication of ZIKV-V153D and ZIKV-N154D in Vero E6 cells and in human brain organoids is recapitulated *in vivo*, these viruses were used to infect *Ifnar^-/-^* mice. Infection with ZIKV-V153D and ZIKV-N154D resulted in the viremia peaking at 1-2 dpi and falling substantially by 4 dpi, whereas the viremia for ZIKV_Natal_ peaked at 3-4 dpi, dropping by 6-7 dpi (Fig. 5A). The glycan mutant viruses thus initially replicated faster, but the viremia was more rapidly curtailed, resulting in a reduced viremic period. ZIKV-V153D and ZIKV-N154D had mean viremia titers >1 log_10_CCID_50_/ml for ∼3.2 days, whereas ZIKV_Natal_ infection showed mean viremia titers >1 log_10_CCID_50_/ml for ∼5 days.

**Figure 5.**
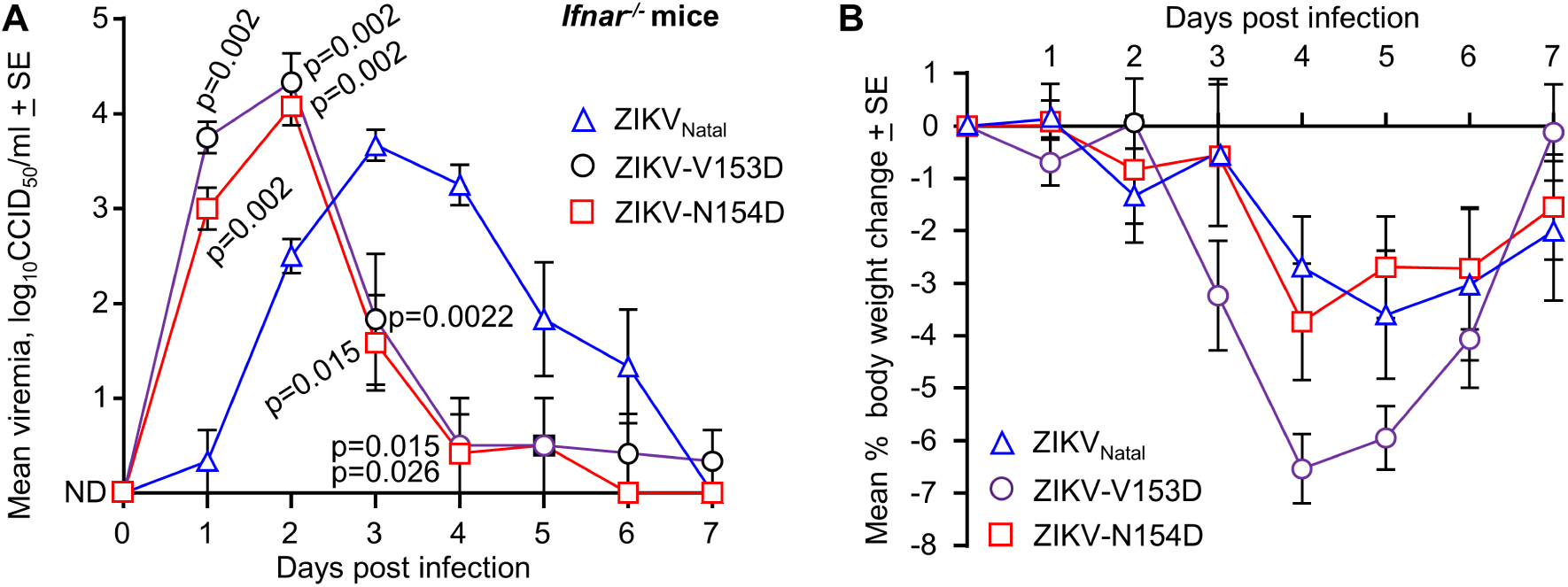
In mice ZIKV-V153D and ZIKV-N154D showed more rapid onset, but shorter viremic periods than ZIKV_Natal_. (A) Mean viremias in *Ifnar^-/-^*mice (n=6 per group) after infection with ZIKV_Natal_, ZIKV-V153D, or ZIKV-N154D. The p values displayed on the graph represent Kolmogorov-Smirnov exact tests, and are all relative to ZIKV_Natal_. At 1 dpi the mean titer for ZIKV-V153D was 0.75 logs higher than for ZIKV-N154D, p = 0.026, t test (this p value is not displayed on the graph). The peak mean titers for ZIKV-V153D and ZIKV-N154D (2 dpi) were ∼0.7 and ∼0.4 logs higher, respectively, than the mean peak titer for ZIKV_Natal_ (3 dpi), although these differences did not reach significance. ND – not detected in all mice; limit of detection per mouse was ∼2 log_10_CCID_50_/ml. (B) Mean percentage change in mouse body weight for the mice described in A.

Spleen was harvested from a subgroup of mice at 3 dpi and the N-linked glycosylation site analyzed by capillary sequencing. The results illustrated that no reversion to intact glycosylation motifs had occurred by 3 dpi for ZIKV-V153D and ZIKV-N154D (Supplementary Fig. 6). Although restoration of the glycosylation motif was observed for ZIKV-MR766-NIID (7), it should be noted that ZIKV-MR766-NIID produces a peak viremia of ∼6 log_10_CCID_50_/ml in *Ifnar^-/-^* mice (7), ∼2 logs higher than those seen herein for ZIKV_Natal_ (Fig. 5A). Less replication provides less opportunity for mutation and selection.

Monitoring of body weight post infection indicated clear reductions in weight, although no mouse reached the ethically defined end point of >15% body weight loss (Fig. 5B). Infection with ZIKV-V153D virus was associated with some increased weight loss compared to the other viruses (Fig. 5B), perhaps due to the slightly higher mean viremia (Fig. 5A). Importantly, infection with the glycosylated parental ZIKV_Natal_ was not associated with increased levels of weight loss or overt pathology when compared to the glycan mutants, arguing against a key role for envelope N-linked glycosylation in driving pathology (18).

## DISCUSSION

We show herein that serial passage of ZIKV_Natal_ in *Rag1*^-/-^ mice selected for viruses with disruptions to the envelope N-linked glycosylation motif. Two of these, ZIKV-V153D and ZIKV-N154D, showed increased replication kinetics in Vero E6 cells and in human brain organoids (up to ∼2.6 logs higher), when compared with the parental glycosylated ZIKV_Natal_. These observations are consistent with studies on other flaviviruses, which show increased replication of envelope N-linked glycan mutants in brains when using animals models where antibodies are largely absent due to young age (26, 27, 98), immunodeficiency (99, 100) or intracranial delivery (13, 25).

Our observations do not support the oft-repeated contention that envelope N-linked glycosylation of ZIKV (and other flaviviruses) is required for proper folding, processing, stability, maturation and/or transport of envelope, and the assembling virion, into the extracellular milieu (19-24). However, such findings may often arise from comparisons of wild-type virus with an engineered glycosylation-motif mutant, where the envelope amino acid sequence changes may impart the key unfavorable properties, with the loss of the glycan residue a secondary phenomenon. For instance, the N154A substitution (35, 39, 40, 101), may itself promote instability, insolubility and/or degradation (22) and/or toxicity (43). The data presented herein argues that the best method for disrupting the glycosylation motif, without introducing a detrimental amino acid change is N154D or V153D. Although these substitutions have previously been identified (37, 102), to the best of our knowledge, N154D or V153D mutants have not been used to study the role of glycosylation (Supplementary Table 1).

ZIKV would not likely have evolved its envelope N-linked glycosylation in order to improve virus replication, as D in position 153 or 154 achieves this goal with substantially greater efficiency. A critical role for lectin-assisted infection processes (17, 53), is also not supported by our data. Instead, our data is consistent with a role for these glycans in the evasion of neutralizing antibody responses. This is not a new concept and has been proposed previously for ZIKV, based on the proximity of the glycan moieties to the fusion loop (15, 16) (Supplementary Fig. 7A) and was recently supported by experimental data (17). Glycan-mediated evasion of neutralizing antibodies has also been reported for other viruses (103-105) and is clearly illustrated by the technique known as “glycan masking” whereby glycans are introduced to specific regions in the immunogen to reduce antibodies response to those regions (61). A key consequence of evading early neutralizing responses is to extend the viremic period, shown herein in Fig. 5A. Although not a selection pressure in play in the current setting, increasing the viremic period would increase the probability of virus transmission from the mammalian host to mosquito vectors, an important factor in arbovirus spread and evolutionary selection (106, 107).

So why do the glycan mutants replicate better? Firstly, RNA-Seq analysis of ZIKV-N154D vs. ZIKV_Natal_ infection of organoids indicated that ZIKV-N154D infection led to increased stimulation of the UPR. Importantly, this was not associated with increased neuron cell death signatures, which were actually slightly reduced. That infection with flaviviruses (including ZIKV) induces the UPR is well documented, with the UPR generally viewed as promoting viral replication (88-90, 108-110). A UPR-mediated upregulation of *inter alia* secretory pathway chaperones and foldases, benefits viral replication, as movement of viral structural proteins and assembling virions through the ER and Golgi and out of the cell is accelerated (95-97). The UPR can lead to apoptosis if the ER stress is not resolved, with UPR-driven apoptosis widely studied (111). However, if stress is resolved, with unfolded proteins cleared from the ER, then cell death can be avoided or delayed (94, 112). In viral infection settings, prolonging host cell survival would clearly promote virus replication (113, 114). A second, likely related, reason why ZIKV-N154D and ZIKV-V153D replicate better may be associated with the high metabolic and kinetic costs of glycosylation (115, 116). Glycosylation involves numerous sequential and competitive steps in an assembly line-like process (117) that requires both energy and time to complete (118). Such costs may be particularly critical during flavivirus replication, where the ER is extensively reorganized to prioritize synthesis and secretion of glycosylated flavivirus structural proteins (20). The bioinformatic analysis suggests increased ER/Golgi transit and viral egress for ZIKV-N154D, arguably this would (i) be easier to achieve without the requirement for glycosylation and (ii) be consistent with a more effective UPR. Thirdly, the V153D and N154D substitutions may not impart unfavorable properties, unlike, for instance, the N154A substitution, which imparts instability, insolubility and degradation (22) and toxicity (43). Perhaps important is that both N154D and V153D preserve (via the aspartic acid residues) a negative charge on the surface of the envelope protein in the location of the glycan loop (Supplementary Fig. 7B). Glycosylated ZIKVs also have a negative charge in this location, provided by the sialic acid residues of the glycan loop (119). Such surface negative charge is lost for the N154A substitution (Supplementary Fig. 7B). One might thus speculate that the N154D and V153D substitutions can replace the glycan moiety, without imparting unfavorable properties, by preserving negative charge at the glycosylation motif site (120). A related observation was reported for *Saccharomyces cerevisiae* proteins expressed under nutrient deficient conditions, which resulted in loss of N-glycosylation and the introduction of charged amino acids at the N-glycosylation sites. These latter amino acid substitutions increased protein stability and activity (121).

Curiously some flaviviruses appear not to carry an envelope N-linked glycan. For instance, Kunjin virus isolates are often not glycosylated; however, this may be due to passage history, rather than a feature of the virus in naturally infected mammalian hosts (122). A primary isolate from a horse did, for instance, have an N-linked envelope glycan (123). Loss of the envelope N-linked glycosylation motif associated with *in vitro* culture has also been reported for a number of other flaviviruses (29, 124, 125). Thus both passage in *Rag1*^-/-^ mice and passage *in vitro* can select viruses devoid of N-linked envelope glycosylation, with both selections occurring under conditions devoid of neutralizing antibodies.

The paper has a number of limitations. Although we have assessed replication in C6/36 cells, we have not assessed fitness in mosquitoes, leaving open the question of whether ZIKV-V153D and ZIKV-N154D are transmitted with the same efficiency as ZIKV_Natal_ (58, 126, 127). Also unclear is why ZIKV-V153I/T156I replicated so poorly in C6/36 cells (Supplementary Fig. 3). Retention of a positively charged patch at the glycan loop site (Supplementary Fig. 7) would not appear to be important for replication in C6/36 cells (Fig. 3B), suggesting the V153I/T156I mutations impart a different disadvantage in insect cells that does not manifest in *Rag1*^-/-^ mice. That replication in insect and mammalian cells selects for quite distinct sets of substitutions in ZIKV_Natal_ envelope has been demonstrated by deep mutational scanning experiments (45). Finally, we are unable to provide cogent insights into the other substitutions outside the glycosylation motif (Fig. 2A, Supplementary Figs. 2C and 4A), except that they are entirely different for each of the three selected glycan mutants (Fig. 2A), none match amino acids seen in the mouse-adapted ZIKV-MR766 strain (Supplementary Table 3), none are located in envelope, and none, to the best of our knowledge, have been associated with increased viral fitness in mammalian cells.

In summary, our data argues that the N-linked glycosylation of the ZIKV envelope protein is not essential for supporting envelope folding, processing, assembly and/or transit through the ER and Golgi. If anything, glycosylation slows this process, with the glycan mutants ZIKV-V153D and ZIKV-N154D producing ∼1-2.6 logs more virus. Instead, N-linked glycosylation of envelope was associated with prolonging the viremia period, consistent with “glycan masking” activity that delays the generation and activity of very early neutralizing antibody responses. One potential application arising from these observations is that yield and immunogenicity of virus-like-particle vaccines and/or yield of viral particle therapeutics (128, 129) might be improved by introduction of either of these D mutations into the glycosylation motif.

## ACKNOWLEDGEMENTS

From QIMR Berghofer MRI we thank: the animal house staff for mouse breeding and agistment; Drs Anthony White and Lotta Oikari for providing the iPSC line; Crystal Chang and Ashwini Potadar for the organoid immunohistochemistry; and Dr. Gunter Hartel for assistance with statistics.

## AUTHOR CONTRIBUTIONS

Conceptualization, A.S. and D.J.R.; methodology: R.S., A.S., A. Sl. and D.J.R.; investigation: B.T., R.S, A.C., K.Y., T.D., W.N., A.K.,; resources: R.S., E.N., A.Sl, A.K.; formal analysis: C.B., D.J.R.; E.N., T.D.; writing-original draft: D.J.R., A.S.; writing-review and editing: A.S. and V.L.; visualization: A.S., D.J.R.; supervision: A.S., D.J.R.; funding acquisition: A.S.

## DATA AVAILABILITY

All data is provided in the manuscript and accompanying supplementary files. Raw sequencing data (fastq files) generated for this publication have been deposited in the NCBI SRA, BioProject: PRJNA1141163 and are publicly available as of the date of publication.

## SUPPLEMENTAL MATERIAL

The manuscript contains four Supplemental Tables and seven Supplementary Figures.

## REFERENCES

1. Saba Villarroel PM, Hamel R, Gumpangseth N, Yainoy S, Koomhin P, Missé D, Wichit S. 2024. Global seroprevalence of Zika virus in asymptomatic individuals: A systematic review. PLOS Neglected Tropical Diseases 18:e0011842. 10.1371/journal.pntd.0011842.

2. Moura LM, Ferreira VLdR, Loureiro RM, de Paiva JPQ, Rosa-Ribeiro R, Amaro E, Soares MBP, Machado BS. 2021. The Neurobiology of Zika Virus: New Models, New Challenges. Frontiers in Neuroscience 15:654078. 10.3389/fnins.2021.654078.

3. Freitas DA, Souza-Santos R, Carvalho LMA, Barros WB, Neves LM, Brasil P, Wakimoto MD. 2020. Congenital Zika syndrome: A systematic review. PLOS ONE 15:e0242367. 10.1371/journal.pone.0242367.

4. Marques FJP, Tran L, Kousa YA, Leyser M. 2024. Long-term developmental outcomes of children with congenital Zika syndrome. Pediatric Research doi:10.1038/s41390-024-03389-9. 10.1038/s41390-024-03389-9.

5. Reynoso GV, Gordon DN, Kalia A, Aguilar CC, Malo CS, Aleshnick M, Dowd KA, Cherry CR, Shannon JP, Vrba SM, Holmes AC, Alippe Y, Maciejewski S, Asano K, Diamond MS, Pierson TC, Hickman HD. 2023. Zika virus spreads through infection of lymph node-resident macrophages. Cell Reports 42:112126. 10.1016/j.celrep.2023.112126.

6. Henrio Marcellin DF, Huang J. 2024. Exploring Zika Virus Impact on Endothelial Permeability: Insights into Transcytosis Mechanisms and Vascular Leakage. Viruses 16:629. 10.3390/v16040629.

7. Nakayama E, Kato F, Tajima S, Ogawa S, Yan K, Takahashi K, Sato Y, Suzuki T, Kawai Y, Inagaki T, Taniguchi S, Le TT, Tang B, Prow NA, Uda A, Maeki T, Lim C-K, Khromykh AA, Suhrbier A, Saijo M. 2021. Neuroinvasiveness of the MR766 strain of Zika virus in IFNAR-/- mice maps to prM residues conserved amongst African genotype viruses. PLOS Pathogens 17:e1009788. 10.1371/journal.ppat.1009788.

8. Komarasamy TV, Adnan NAA, James W, Balasubramaniam VRMT. 2022. Zika Virus Neuropathogenesis: The Different Brain Cells, Host Factors and Mechanisms Involved. Frontiers in Immunology 13:773191. 10.3389/fimmu.2022.773191.

9. Nakayama E, Kawai Y, Taniguchi S, Hazlewood JE, Shibasaki K-i, Takahashi K, Sato Y, Tang B, Yan K, Katsuta N, Tajima S, Lim CK, Suzuki T, Suhrbier A, Saijo M. 2021. Embryonic Stage of Congenital Zika Virus Infection Determines Fetal and Postnatal Outcomes in Mice. Viruses 13:1807. 10.3390/v13091807.

10. Goo L, VanBlargan LA, Dowd KA, Diamond MS, Pierson TC. 2017. A single mutation in the envelope protein modulates flavivirus antigenicity, stability, and pathogenesis. PLOS Pathogens 13:e1006178. 10.1371/journal.ppat.1006178.

11. Fontes-Garfias CR, Shan C, Luo H, Muruato AE, Medeiros DBA, Mays E, Xie X, Zou J, Roundy CM, Wakamiya M, Rossi SL, Wang T, Weaver SC, Shi PY. 2017. Functional Analysis of Glycosylation of Zika Virus Envelope Protein. Cell Rep 21:1180–1190. 10.1016/j.celrep.2017.10.016.

12. Giraldo MI, Xia H, Aguilera-Aguirre L, Hage A, van Tol S, Shan C, Xie X, Sturdevant GL, Robertson SJ, McNally KL, Meade-White K, Azar SR, Rossi SL, Maury W, Woodson M, Ramage H, Johnson JR, Krogan NJ, Morais MC, Best SM, Shi P-Y, Rajsbaum R. 2020. Envelope protein ubiquitination drives entry and pathogenesis of Zika virus. Nature 585:414–419. 10.1038/s41586-020-2457-8.

13. Cheng M-L, Yang Y-X, Liu Z-Y, Wen D, Yang P, Huang X-Y, Dong H-L, Xu Y-P, Li X-F, Deng Y-Q, Ye Q, Zhu L, Li J, Davidson Andrew D, Zheng A-H, Shi W-F, Zhao H, Wang X-X, Qin C-F. 2022. Pathogenicity and Structural Basis of Zika Variants with Glycan Loop Deletions in the Envelope Protein. Journal of Virology 96:e00879–22. 10.1128/jvi.00879-22.

14. Rawle DJ, Hugo LE, Cox AL, Devine GJ, Suhrbier A. 2024. Generating prophylactic immunity against arboviruses in vertebrates and invertebrates. Nat Rev Immunol 24:621–636. 10.1038/s41577-024-01016-6.

15. Sirohi D, Chen Z, Sun L, Klose T, Pierson TC, Rossmann MG, Kuhn RJ. 2016. The 3.8 Å resolution cryo-EM structure of Zika virus. Science 352:467–470. 10.1126/science.aaf5316.

16. Frumence E, Viranaicken W, Bos S, Alvarez-Martinez M-T, Roche M, Arnaud J-D, Gadea G, Desprès P. 2019. A Chimeric Zika Virus between Viral Strains MR766 and BeH819015 Highlights a Role for E-glycan Loop in Antibody-mediated Virus Neutralization. Vaccines 7:55. 10.3390/vaccines7020055.

17. Contreras M, Stuart JB, Levoir LM, Belmont L, Goo L. 2024. Defining the impact of flavivirus envelope protein glycosylation site mutations on sensitivity to broadly neutralizing antibodies. mBio 15:e0304823. 10.1128/mbio.03048-23.

18. Feng T, Zhang J, Chen Z, Pan W, Chen Z, Yan Y, Dai J. 2022. Glycosylation of viral proteins: Implication in virus–host interaction and virulence. Virulence 13:670–683. 10.1080/21505594.2022.2060464.

19. Courageot M-P, Frenkiel M-P, Duarte Dos Santos C, Deubel V, Desprès P. 2000. α-Glucosidase Inhibitors Reduce Dengue Virus Production by Affecting the Initial Steps of Virion Morphogenesis in the Endoplasmic Reticulum. Journal of Virology 74:564-572. 10.1128/jvi.74.1.564-572.2000.

20. Verhaegen M, Vermeire K. 2024. The endoplasmic reticulum (ER): a crucial cellular hub in flavivirus infection and potential target site for antiviral interventions. npj Viruses 2:24. 10.1038/s44298-024-00031-7.

21. Pandey B, S S, Chatterjee A, Mangala Prasad V. 2023. Role of surface glycans in enveloped RNA virus infections: A structural perspective. Proteins doi:10.1002/prot.26636:1-12. 10.1002/prot.26636.

22. Ishida K, Yagi H, Kato Y, Morita E. 2023. N-linked glycosylation of flavivirus E protein contributes to viral particle formation. PLOS Pathogens 19:e1011681. 10.1371/journal.ppat.1011681.

23. Mossenta M, Marchese S, Poggianella M, Slon Campos JL, Burrone OR. 2017. Role of N-glycosylation on Zika virus E protein secretion, viral assembly and infectivity. Biochemical and Biophysical Research Communications 492:579–586. 10.1016/j.bbrc.2017.01.022.

24. Gwon Y-D, Zusinaite E, Merits A, Överby AK, Evander M. 2020. N-glycosylation in the Pre-Membrane Protein Is Essential for the Zika Virus Life Cycle. Viruses 12:925. 10.3390/v12090925.

25. Fang E, Liu X, Li M, Liu J, Zhang Z, Liu X, Li X, Li W, Peng Q, Yu Y, Li Y. 2022. Construction of a Dengue NanoLuc Reporter Virus for In Vivo Live Imaging in Mice. Viruses 14:1253. 10.3390/v14061253.

26. Guo Y, Bao L, Xu Y, Li F, Lv Q, Fan F, Qin C. 2021. The Ablation of Envelope Protein Glycosylation Enhances the Neurovirulence of ZIKV and Cell Apoptosis in Newborn Mice. Journal of Immunology Research 2021:5317662. 10.1155/2021/5317662.

27. Kariwa H, Murata R, Totani M, Yoshii K, Takashima I. 2013. Increased pathogenicity of West Nile virus (WNV) by glycosylation of envelope protein and seroprevalence of WNV in wild birds in Far Eastern Russia. Int J Environ Res Public Health 10:7144–64. 10.3390/ijerph10127144.

28. Chambers TJ, Halevy M, Nestorowicz A, Rice CM, Lustig S. 1998. West Nile virus envelope proteins: nucleotide sequence analysis of strains differing in mouse neuroinvasiveness. J Gen Virol 79 (Pt 10):2375–80. 10.1099/0022-1317-79-10-2375.

29. Lee E, Weir RC, Dalgarno L. 1997. Changes in the dengue virus major envelope protein on passaging and their localization on the three-dimensional structure of the protein. Virology 232:281–90. 10.1006/viro.1997.8570.

30. Zhang Y, Yan Y, Li S, Yuan F, Wen D, Jia N, Xiong T, Zhang X, Zheng A. 2023. Broad Host Tropism of Flaviviruses during the Entry Stage. Microbiology Spectrum 11:e05281–22. 10.1128/spectrum.05281-22.

31. Maharaj PD, Langevin SA, Bolling BG, Andrade CC, Engle XA, Ramey WN, Bosco-Lauth A, Bowen RA, Sanders TA, Huang CYH, Reisen WK, Brault AC. 2019. N-linked glycosylation of the West Nile virus envelope protein is not a requisite for avian virulence or vector competence. PLOS Neglected Tropical Diseases 13:e0007473. 10.1371/journal.pntd.0007473.

32. Alen MM, Dallmeier K, Balzarini J, Neyts J, Schols D. 2012. Crucial role of the N-glycans on the viral E-envelope glycoprotein in DC-SIGN-mediated dengue virus infection. Antiviral Res 96:280–7. 10.1016/j.antiviral.2012.10.007.

33. Moudy RM, Payne AF, Dodson BL, Kramer LD. 2011. Requirement of glycosylation of West Nile virus envelope protein for infection of, but not spread within, Culex quinquefasciatus mosquito vectors. Am J Trop Med Hyg 85:374–8. 10.4269/ajtmh.2011.10-0697.

34. Moudy RM, Zhang B, Shi PY, Kramer LD. 2009. West Nile virus envelope protein glycosylation is required for efficient viral transmission by Culex vectors. Virology 387:222–8. 10.1016/j.virol.2009.01.038.

35. Annamalai AS, Pattnaik A, Sahoo B, R., Muthukrishnan E, Natarajan SK, Steffen D, Vu H, Delhon G, Osorio FA, Petro TM, Xiang S-H, Pattnaik AK. 2017. Zika Virus Encoding Nonglycosylated Envelope Protein Is Attenuated and Defective in Neuroinvasion. Journal of Virology 91:e01348–17. 10.1128/jvi.01348-17.

36. Bos S, Viranaicken W, Frumence E, Li G, Despres P, Zhao RY, Gadea G. 2019. The Envelope Residues E152/156/158 of Zika Virus Influence the Early Stages of Virus Infection in Human Cells. Cells 8:1444. 10.3390/cells8111444.

37. Gallichotte EN, Samaras D, Murrieta RA, Sexton NR, Robison A, Young MC, Byas AD, Ebel GD, Ruckert C. 2023. The Incompetence of Mosquitoes-Can Zika Virus Be Adapted To Infect Culex tarsalis Cells? mSphere 8:e0001523. 10.1128/msphere.00015-23.

38. Gong D, Zhang TH, Zhao D, Du Y, Chapa TJ, Shi Y, Wang L, Contreras D, Zeng G, Shi PY, Wu TT, Arumugaswami V, Sun R. 2018. High-Throughput Fitness Profiling of Zika Virus E Protein Reveals Different Roles for Glycosylation during Infection of Mammalian and Mosquito Cells. iScience 1:97–111. 10.1016/j.isci.2018.02.005.

39. Liu D, Xiao X, Zhou P, Zheng H, Li Y, Jin H, Jongkaewwattana A, Luo R. 2021. Glycosylation on envelope glycoprotein of duck Tembusu virus affects virus replication in vitro and contributes to the neurovirulence and pathogenicity in vivo. Virulence 12:2400–2414. 10.1080/21505594.2021.1974329.

40. Liang J-J, Chou M-W, Lin Y-L. 2018. DC-SIGN Binding Contributed by an Extra N-Linked Glycosylation on Japanese Encephalitis Virus Envelope Protein Reduces the Ability of Viral Brain Invasion. Frontiers in Cellular and Infection Microbiology 8:239. 10.3389/fcimb.2018.00239.

41. Prow NA, May FJ, Westlake DJ, Hurrelbrink RJ, Biron RM, Leung JY, McMinn PC, Clark DC, Mackenzie JS, Lobigs M, Khromykh AA, Hall RA. 2011. Determinants of attenuation in the envelope protein of the flavivirus Alfuy. J Gen Virol 92:2286–2296. 10.1099/vir.0.034793-0.

42. Widman DG, Young E, Yount BL, Plante KS, Gallichotte EN, Carbaugh DL, Peck KM, Plante J, Swanstrom J, Heise MT, Lazear HM, Baric RS. 2017. A Reverse Genetics Platform That Spans the Zika Virus Family Tree. mBio 8:10.1128/mbio.02014-16. 10.1128/mBio.02014-16.

43. Steiner JP, Bachani M, Malik N, Li W, Tyagi R, Sampson K, Abrams RPM, Kousa Y, Solis J, Johnson TP, Nath A. 2023. Neurotoxic properties of the Zika virus envelope protein. Exp Neurol 367:114469. 10.1016/j.expneurol.2023.114469.

44. Lazear HM, Govero J, Smith AM, Platt DJ, Fernandez E, Miner JJ, Diamond MS. 2016. A Mouse Model of Zika Virus Pathogenesis. Cell Host Microbe 19:720–30. 10.1016/j.chom.2016.03.010.

45. Setoh YX, Amarilla AA, Peng NYG, Griffiths RE, Carrera J, Freney ME, Nakayama E, Ogawa S, Watterson D, Modhiran N, Nanyonga FE, Torres FJ, Slonchak A, Periasamy P, Prow NA, Tang B, Harrison J, Hobson-Peters J, Cuddihy T, Cooper-White J, Hall RA, Young PR, Mackenzie JM, Wolvetang E, Bloom JD, Suhrbier A, Khromykh AA. 2019. Determinants of Zika virus host tropism uncovered by deep mutational scanning. Nature Microbiology 4:876–887. 10.1038/s41564-019-0399-4.

46. Prow NA, Liu L, Nakayama E, Cooper TH, Yan K, Eldi P, Hazlewood JE, Tang B, Le TT, Setoh YX, Khromykh AA, Hobson-Peters J, Diener KR, Howley PM, Hayball JD, Suhrbier A. 2018. A vaccinia-based single vector construct multi-pathogen vaccine protects against both Zika and chikungunya viruses. Nat Commun 9:1230. 10.1038/s41467-018-03662-6.

47. Hazlewood JE, Rawle DJ, Tang B, Yan K, Vet LJ, Nakayama E, Hobson-Peters J, Hall RA, Suhrbier A. 2020. A Zika Vaccine Generated Using the Chimeric Insect-Specific Binjari Virus Platform Protects against Fetal Brain Infection in Pregnant Mice. Vaccines (Basel) 8:496. 10.3390/vaccines8030496.

48. Calvert AE, Horiuchi K, Boroughs KL, Ong YT, Anderson KM, Biggerstaff BJ, Stone M, Simmons G, Busch MP, Huang CY. 2021. The Specificity of the Persistent IgM Neutralizing Antibody Response in Zika Virus Infections among Individuals with Prior Dengue Virus Exposure. J Clin Microbiol 59:e0040021. 10.1128/jcm.00400-21.

49. Singh T, Hwang KK, Miller AS, Jones RL, Lopez CA, Dulson SJ, Giuberti C, Gladden MA, Miller I, Webster HS, Eudailey JA, Luo K, Von Holle T, Edwards RJ, Valencia S, Burgomaster KE, Zhang S, Mangold JF, Tu JJ, Dennis M, Alam SM, Premkumar L, Dietze R, Pierson TC, Ooi EE, Lazear HM, Kuhn RJ, Permar SR, Bonsignori M. 2022. A Zika virus-specific IgM elicited in pregnancy exhibits ultrapotent neutralization. Cell 185:4826–4840 e17. 10.1016/j.cell.2022.10.023.

50. Lawson CM, Grundy JE, Shellam GR. 1988. Antibody responses to murine cytomegalovirus in genetically resistant and susceptible strains of mice. J Gen Virol 69 (Pt 8):1987–98. 10.1099/0022-1317-69-8-1987.

51. Zhang Y-Y, Summers J. 2004. Rapid Production of Neutralizing Antibody Leads to Transient Hepadnavirus Infection. Journal of Virology 78:1195–1201. 10.1128/jvi.78.3.1195-1201.2004.

52. Al-Ibraheemi JSS, Al-Saeedi AS. 2021. The relationship between IgG and IgM levels and severity of symptoms in COVID-19 patients confirmed by rapid antigen test. J Med Life 14:790–796. 10.25122/jml-2021-0194.

53. Carbaugh DL, Lazear HM. 2020. Flavivirus Envelope Protein Glycosylation: Impacts on Viral Infection and Pathogenesis. J Virol 94. 10.1128/jvi.00104-20.

54. Gorman MJ, Caine EA, Zaitsev K, Begley MC, Weger-Lucarelli J, Uccellini MB, Tripathi S, Morrison J, Yount BL, Dinnon KH, Rückert C, Young MC, Zhu Z, Robertson SJ, McNally KL, Ye J, Cao B, Mysorekar IU, Ebel GD, Baric RS, Best SM, Artyomov MN, Garcia-Sastre A, Diamond MS. 2018. An Immunocompetent Mouse Model of Zika Virus Infection. Cell Host & Microbe 23:672–685.e6. 10.1016/j.chom.2018.04.003.

55. Grant A, Ponia SS, Tripathi S, Balasubramaniam V, Miorin L, Sourisseau M, Schwarz MC, Sanchez-Seco MP, Evans MJ, Best SM, Garcia-Sastre A. 2016. Zika Virus Targets Human STAT2 to Inhibit Type I Interferon Signaling. Cell Host Microbe 19:882–90. 10.1016/j.chom.2016.05.009.

56. Kumar A, Hou S, Airo AM, Limonta D, Mancinelli V, Branton W, Power C, Hobman TC. 2016. Zika virus inhibits type-I interferon production and downstream signaling. EMBO Rep 17:1766–1775. 10.15252/embr.201642627.

57. Magnani DM, Rogers TF, Beutler N, Ricciardi MJ, Bailey VK, Gonzalez-Nieto L, Briney B, Sok D, Le K, Strubel A, Gutman MJ, Pedreño-Lopez N, Grubaugh ND, Silveira CGT, Maxwell HS, Domingues A, Martins MA, Lee DE, Okwuazi EE, Jean S, Strobert EA, Chahroudi A, Silvestri G, Vanderford TH, Kallas EG, Desrosiers RC, Bonaldo MC, Whitehead SS, Burton DR, Watkins DI. 2017. Neutralizing human monoclonal antibodies prevent Zika virus infection in macaques. Science Translational Medicine 9:eaan8184. 10.1126/scitranslmed.aan8184.

58. Setoh YX, Prow NA, Peng N, Hugo LE, Devine G, Hazlewood JE, Suhrbier A, Khromykh AA. 2017. De Novo Generation and Characterization of New Zika Virus Isolate Using Sequence Data from a Microcephaly Case. mSphere 2:e00190–17. 10.1128/mSphereDirect.00190-17.

59. Mlakar J, Korva M, Tul N, Popović M, Poljšak-Prijatelj M, Mraz J, Kolenc M, Resman Rus K, Vesnaver Vipotnik T, Fabjan Vodušek V, Vizjak A, Pižem J, Petrovec M, Avšič Županc T. 2016. Zika Virus Associated with Microcephaly. N Engl J Med 374:951–8. 10.1056/NEJMoa1600651.

60. Shao Q, Herrlinger S, Zhu Y-N, Yang M, Goodfellow F, Stice SL, Qi X-P, Brindley MA, Chen J-F. 2017. The African Zika virus MR-766 is more virulent and causes more severe brain damage than current Asian lineage and dengue virus. Development 144:4114–4124. 10.1242/dev.156752.

61. Martina CE, Crowe JE, Jr., Meiler J. 2023. Glycan masking in vaccine design: Targets, immunogens and applications. Front Immunol 14:1126034. 10.3389/fimmu.2023.1126034.

62. Sanders RW, van Anken E, Nabatov AA, Liscaljet IM, Bontjer I, Eggink D, Melchers M, Busser E, Dankers MM, Groot F, Braakman I, Berkhout B, Paxton WA. 2008. The carbohydrate at asparagine 386 on HIV-1 gp120 is not essential for protein folding and function but is involved in immune evasion. Retrovirology 5:10. 10.1186/1742-4690-5-10.

63. Yan K, Dumenil T, Tang B, Le TT, Bishop CR, Suhrbier A, Rawle DJ. 2022. Evolution of ACE2-independent SARS-CoV-2 infection and mouse adaption after passage in cells expressing human and mouse ACE2. Virus Evol 8:veac063. 10.1093/ve/veac063.

64. Anonymous. TCID50_calculator_v2_17-01-20_MB - Excel sheet to calculate TCID_50_ titers (Spearman & Kärber method). https://www.klinikum.uni-heidelberg.de/zentrum-fuer-infektiologie/molecular-virology/welcome/downloads. Accessed Feb 2024.

65. Slonchak A, Wang X, Aguado J, Sng JDJ, Chaggar H, Freney ME, Yan K, Torres FJ, Amarilla AA, Balea R, Setoh YX, Peng N, Watterson D, Wolvetang E, Suhrbier A, Khromykh AA. 2022. Zika virus noncoding RNA cooperates with the viral protein NS5 to inhibit STAT1 phosphorylation and facilitate viral pathogenesis. Sci Adv 8:eadd8095. 10.1126/sciadv.add8095.

66. Oikari LE, Pandit R, Stewart R, Cuni-Lopez C, Quek H, Sutharsan R, Rantanen LM, Oksanen M, Lehtonen S, de Boer CM, Polo JM, Gotz J, Koistinaho J, White AR. 2020. Altered Brain Endothelial Cell Phenotype from a Familial Alzheimer Mutation and Its Potential Implications for Amyloid Clearance and Drug Delivery. Stem Cell Reports 14:924–939. 10.1016/j.stemcr.2020.03.011.

67. Stewart R, Yan K, Ellis SA, Bishop CR, Dumenil T, Tang B, Nguyen W, Larcher T, Parry R, Sng JDJ, Khromykh AA, Sullivan RKP, Lor M, Meunier FA, Rawle DJ, Suhrbier A. 2023. SARS-CoV-2 omicron BA.5 and XBB variants have increased neurotropic potential over BA.1 in K18-hACE2 mice and human brain organoids. Frontiers in Microbiology 14:1320856. 10.3389/fmicb.2023.1320856.

68. Nguyen W, Gyawali N, Stewart R, Tang B, Cox AL, Yan K, Larcher T, Bishop CR, Wood N, Devine GJ, Suhrbier A, Rawle DJ. 2024. Characterisation of a Japanese Encephalitis virus genotype 4 isolate from the 2022 Australian outbreak. npj Viruses 2:15. 10.1038/s44298-024-00025-5.

69. Nguyen W, Nakayama E, Yan K, Tang B, Le TT, Liu L, Cooper TH, Hayball JD, Faddy HM, Warrilow D, Allcock RJN, Hobson-Peters J, Hall RA, Rawle DJ, Lutzky VP, Young P, Oliveira NM, Hartel G, Howley PM, Prow NA, Suhrbier A. 2020. Arthritogenic Alphavirus Vaccines: Serogrouping Versus Cross-Protection in Mouse Models. Vaccines 8:209. 10.3390/vaccines8020209.

70. Schneider CA, Rasband WS, Eliceiri KW. 2012. NIH Image to ImageJ: 25 years of image analysis. Nat Methods 9:671–5. 10.1038/nmeth.2089.

71. Hazlewood JE, Dumenil T, Le TT, Slonchak A, Kazakoff SH, Patch AM, Gray LA, Howley PM, Liu L, Hayball JD, Yan K, Rawle DJ, Prow NA, Suhrbier A. 2021. Injection site vaccinology of a recombinant vaccinia-based vector reveals diverse innate immune signatures. PLoS Pathog 17:e1009215. 10.1371/journal.ppat.1009215.

72. Langmead B, Salzberg SL. 2012. Fast gapped-read alignment with Bowtie 2. Nat Methods 9:357–9. 10.1038/nmeth.1923.

73. Li H, Handsaker B, Wysoker A, Fennell T, Ruan J, Homer N, Marth G, Abecasis G, Durbin R. 2009. The Sequence Alignment/Map format and SAMtools. Bioinformatics 25:2078–9. 10.1093/bioinformatics/btp352.

74. Dobin A, Davis CA, Schlesinger F, Drenkow J, Zaleski C, Jha S, Batut P, Chaisson M, Gingeras TR. 2013. STAR: ultrafast universal RNA-seq aligner. Bioinformatics 29:15–21. 10.1093/bioinformatics/bts635.

75. Li B, Dewey CN. 2011. RSEM: accurate transcript quantification from RNA-Seq data with or without a reference genome. BMC Bioinformatics 12:323. 10.1186/1471-2105-12-323.

76. Love MI, Huber W, Anders S. 2014. Moderated estimation of fold change and dispersion for RNA-seq data with DESeq2. Genome Biol 15:550. 10.1186/s13059-014-0550-8.

77. Subramanian A, Tamayo P, Mootha VK, Mukherjee S, Ebert BL, Gillette MA, Paulovich A, Pomeroy SL, Golub TR, Lander ES, Mesirov JP. 2005. Gene set enrichment analysis: a knowledge-based approach for interpreting genome-wide expression profiles. Proc Natl Acad Sci U S A 102:15545–50. 10.1073/pnas.0506580102.

78. Wilm A, Aw PP, Bertrand D, Yeo GH, Ong SH, Wong CH, Khor CC, Petric R, Hibberd ML, Nagarajan N. 2012. LoFreq: a sequence-quality aware, ultra-sensitive variant caller for uncovering cell-population heterogeneity from high-throughput sequencing datasets. Nucleic Acids Res 40:11189–201. 10.1093/nar/gks918.

79. Van der Auwera G, O’Connor B. 2020. Genomics in the Cloud: Using Docker, GATK, and WDL in Terra, 1 ed. O’Reilly Media, California, United States.

80. Swann JB, Hayakawa Y, Zerafa N, Sheehan KC, Scott B, Schreiber RD, Hertzog P, Smyth MJ. 2007. Type I IFN contributes to NK cell homeostasis, activation, and antitumor function. J Immunol 178:7540–9. 10.4049/jimmunol.178.12.7540.

81. Rawle DJ, Le TT, Dumenil T, Bishop C, Yan K, Nakayama E, Bird PI, Suhrbier A. 2022. Widespread discrepancy in Nnt genotypes and genetic backgrounds complicates granzyme A and other knockout mouse studies. Elife 11:e70207. 10.7554/eLife.70207.

82. Poo YS, Rudd PA, Gardner J, Wilson JA, Larcher T, Colle MA, Le TT, Nakaya HI, Warrilow D, Allcock R, Bielefeldt-Ohmann H, Schroder WA, Khromykh AA, Lopez JA, Suhrbier A. 2014. Multiple immune factors are involved in controlling acute and chronic chikungunya virus infection. PLoS Negl Trop Dis 8:e3354. 10.1371/journal.pntd.0003354.

83. Hayashida E, Ling ZL, Ashhurst TM, Viengkhou B, Jung SR, Songkhunawej P, West PK, King NJC, Hofer MJ. 2019. Zika virus encephalitis in immunocompetent mice is dominated by innate immune cells and does not require T or B cells. Journal of Neuroinflammation 16:177. 10.1186/s12974-019-1566-5.

84. Sutarjono B. 2019. Can We Better Understand How Zika Leads to Microcephaly? A Systematic Review of the Effects of the Zika Virus on Human Brain Organoids. J Infect Dis 219:734–745. 10.1093/infdis/jiy572.

85. Passemard S, Perez F, Gressens P, El Ghouzzi V. 2019. Endoplasmic reticulum and Golgi stress in microcephaly. Cell Stress 3:369–384. 10.15698/cst2019.12.206.

86. Liu S, Costa M. 2022. The role of NUPR1 in response to stress and cancer development. Toxicology and Applied Pharmacology 454:116244. 10.1016/j.taap.2022.116244.

87. Neill G, Masson GR. 2023. A stay of execution: ATF4 regulation and potential outcomes for the integrated stress response. Frontiers in Molecular Neuroscience 16:1112253. 10.3389/fnmol.2023.1112253.

88. Kolpikova EP, Tronco AR, Den Hartigh AB, Jackson KJ, Iwawaki T, Fink SL. 2020. IRE1α Promotes Zika Virus Infection via XBP1. Viruses 12:278. 10.3390/v12030278.

89. Breitkopf VJM, Dobler G, Claus P, Naim HY, Steffen I. 2021. IRE1-Mediated Unfolded Protein Response Promotes the Replication of Tick-Borne Flaviviruses in a Virus and Cell-Type Dependent Manner. Viruses 13:2164. 10.3390/v13112164.

90. Tan Z, Zhang W, Sun J, Fu Z, Ke X, Zheng C, Zhang Y, Li P, Liu Y, Hu Q, Wang H, Zheng Z. 2018. ZIKV infection activates the IRE1-XBP1 and ATF6 pathways of unfolded protein response in neural cells. J Neuroinflammation 15:275. 10.1186/s12974-018-1311-5.

91. Kim E-K, Kim Y, Yang JY, Jang HH. 2022. Prx1 Regulates Thapsigargin-Mediated UPR Activation and Apoptosis. Genes 13:2033. 10.3390/genes13112033.

92. Jiao B, Zhang M, Zhang C, Cao X, Liu B, Li N, Sun J, Zhang X. 2023. Transcriptomics reveals the effects of NTRK1 on endoplasmic reticulum stress response-associated genes in human neuronal cell lines. PeerJ 11:e15219. 10.7717/peerj.15219.

93. Ajoolabady A, Wang S, Kroemer G, Klionsky DJ, Uversky VN, Sowers JR, Aslkhodapasandhokmabad H, Bi Y, Ge J, Ren J. 2021. ER Stress in Cardiometabolic Diseases: From Molecular Mechanisms to Therapeutics. Endocrine Reviews 42:839–871. 10.1210/endrev/bnab006.

94. Read A, Schröder M. 2021. The Unfolded Protein Response: An Overview. Biology 10:384. 10.3390/biology10050384.

95. Shaheen A. 2018. Effect of the unfolded protein response on ER protein export: a potential new mechanism to relieve ER stress. Cell Stress and Chaperones 23:797–806. 10.1007/s12192-018-0905-2.

96. Raschmanová H, Weninger A, Knejzlík Z, Melzoch K, Kovar K. 2021. Engineering of the unfolded protein response pathway in Pichia pastoris: enhancing production of secreted recombinant proteins. Applied Microbiology and Biotechnology 105:4397–4414. 10.1007/s00253-021-11336-5.

97. Lin Y, Feng Y, Zheng L, Zhao M, Huang M. 2023. Improved protein production in yeast using cell engineering with genes related to a key factor in the unfolded protein response. Metabolic Engineering 77:152–161. 10.1016/j.ymben.2023.04.004.

98. Kawano H, Rostapshov V, Rosen L, Lai CJ. 1993. Genetic determinants of dengue type 4 virus neurovirulence for mice. J Virol 67:6567–75. 10.1128/JVI.67.11.6567-6575.1993.

99. Halevy M, Akov Y, Ben-Nathan D, Kobiler D, Lachmi B, Lustig S. 1994. Loss of active neuroinvasiveness in attenuated strains of West Nile virus: pathogenicity in immunocompetent and SCID mice. Arch Virol 137:355–70. 10.1007/BF01309481.

100. Chambers TJ, Droll DA, Walton AH, Schwartz J, Wold WSM, Nickells J. 2008. West Nile 25A virus infection of B-cell-deficient ((micro)MT) mice: characterization of neuroinvasiveness and pseudoreversion of the viral envelope protein. J Gen Virol 89:627–635. 10.1099/vir.0.83297-0.

101. Xu B, Liu X, Yan D, Teng Q, Yuan C, Zhang Z, Liu Q, Li Z. 2023. Generation and characterization of chimeric Tembusu viruses containing pre-membrane and envelope genes of Japanese encephalitis virus. Frontiers in Microbiology 14:1140141. 10.3389/fmicb.2023.1140141.

102. Liu Z, Zhang Y, Cheng M, Ge N, Shu J, Xu Z, Su X, Kou Z, Tong Y, Qin C, Jin X. 2022. A single nonsynonymous mutation on ZIKV E protein-coding sequences leads to markedly increased neurovirulence in vivo. Virologica Sinica 37:115–126. 10.1016/j.virs.2022.01.021.

103. Sommerstein R, Flatz L, Remy MM, Malinge P, Magistrelli G, Fischer N, Sahin M, Bergthaler A, Igonet S, ter Meulen J, Rigo D, Meda P, Rabah N, Coutard B, Bowden TA, Lambert P-H, Siegrist C-A, Pinschewer DD. 2015. Arenavirus Glycan Shield Promotes Neutralizing Antibody Evasion and Protracted Infection. PLOS Pathogens 11:e1005276. 10.1371/journal.ppat.1005276.

104. Karlsson Hedestam GB, Fouchier RAM, Phogat S, Burton DR, Sodroski J, Wyatt RT. 2008. The challenges of eliciting neutralizing antibodies to HIV-1 and to influenza virus. Nature Reviews Microbiology 6:143–155. 10.1038/nrmicro1819.

105. Lavie M, Hanoulle X, Dubuisson J. 2018. Glycan Shielding and Modulation of Hepatitis C Virus Neutralizing Antibodies. Front Immunol 9:910. 10.3389/fimmu.2018.00910.

106. Armstrong PM, Ehrlich HY, Magalhaes T, Miller MR, Conway PJ, Bransfield A, Misencik MJ, Gloria-Soria A, Warren JL, Andreadis TG, Shepard JJ, Foy BD, Pitzer VE, Brackney DE. 2020. Successive blood meals enhance virus dissemination within mosquitoes and increase transmission potential. Nature Microbiology 5:239–247. 10.1038/s41564-019-0619-y.

107. Fesce E, Marini G, Rosa R, Lelli D, Cerioli MP, Chiari M, Farioli M, Ferrari N. 2023. Understanding West Nile virus transmission: Mathematical modelling to quantify the most critical parameters to predict infection dynamics. PLoS Negl Trop Dis 17:e0010252. 10.1371/journal.pntd.0010252.

108. Lewy TG, Grabowski JM, Bloom ME. 2017. BiP: Master Regulator of the Unfolded Protein Response and Crucial Factor in Flavivirus Biology Yale J Biol Med 90:291–300.

109. Yeh CT, Weng SC, Tsao PN, Shiao SH. 2023. The chaperone BiP promotes dengue virus replication and mosquito vitellogenesis in Aedes aegypti. Insect Biochem Mol Biol 155:103930. 10.1016/j.ibmb.2023.103930.

110. Viettri M, Zambrano JL, Rosales R, Caraballo GI, Gutiérrez-Escolano AL, Ludert JE. 2021. Flavivirus infections induce a Golgi stress response in vertebrate and mosquito cells. Scientific Reports 11:23489. 10.1038/s41598-021-02929-1.

111. Kapuy O. 2024. Mechanism of Decision Making between Autophagy and Apoptosis Induction upon Endoplasmic Reticulum Stress. International Journal of Molecular Sciences 25:4368. 10.3390/ijms25084368.

112. Kovaleva V, Yu L-Y, Ivanova L, Shpironok O, Nam J, Eesmaa A, Kumpula E-P, Sakson S, Toots U, Ustav M, Huiskonen JT, Voutilainen MH, Lindholm P, Karelson M, Saarma M. 2023. MANF regulates neuronal survival and UPR through its ER-located receptor IRE1α. Cell Reports 42:112066. 10.1016/j.celrep.2023.112066.

113. Prasad V, Greber UF. 2021. The endoplasmic reticulum unfolded protein response - homeostasis, cell death and evolution in virus infections. FEMS Microbiol Rev 45:fuab016. 10.1093/femsre/fuab016.

114. Ambrose Rebecca L, Mackenzie Jason M. 2011. West Nile Virus Differentially Modulates the Unfolded Protein Response To Facilitate Replication and Immune Evasion. Journal of Virology 85:2723–2732. 10.1128/jvi.02050-10.

115. Mariño K, Bones J, Kattla JJ, Rudd PM. 2010. A systematic approach to protein glycosylation analysis: a path through the maze. Nature Chemical Biology 6:713–723. 10.1038/nchembio.437.

116. del Val IJ, Polizzi KM, Kontoravdi C. 2016. A theoretical estimate for nucleotide sugar demand towards Chinese Hamster Ovary cellular glycosylation. Scientific Reports 6:28547. 10.1038/srep28547.

117. Breitling J, Aebi M. 2013. N-linked protein glycosylation in the endoplasmic reticulum. Cold Spring Harb Perspect Biol 5:a013359. 10.1101/cshperspect.a013359.

118. Scheper AF, Schofield J, Bohara R, Ritter T, Pandit A. 2023. Understanding glycosylation: Regulation through the metabolic flux of precursor pathways. Biotechnology Advances 67:108184. 10.1016/j.biotechadv.2023.108184.

119. Routhu NK, Lehoux SD, Rouse EA, Bidokhti MRM, Giron LB, Anzurez A, Reid SP, Abdel-Mohsen M, Cummings RD, Byrareddy SN. 2019. Glycosylation of Zika Virus is Important in Host–Virus Interaction and Pathogenic Potential. International Journal of Molecular Sciences 20:5206. 10.3390/ijms20205206.

120. Qing R, Hao S, Smorodina E, Jin D, Zalevsky A, Zhang S. 2022. Protein Design: From the Aspect of Water Solubility and Stability. Chemical Reviews 122:14085–14179. 10.1021/acs.chemrev.1c00757.

121. Tan NY, Bailey UM, Jamaluddin MF, Mahmud SH, Raman SC, Schulz BL. 2014. Sequence-based protein stabilization in the absence of glycosylation. Nat Commun 5:3099. 10.1038/ncomms4099.

122. Adams SC, Broom AK, Sammels LM, Hartnett AC, Howard MJ, Coelen RJ, Mackenzie JS, Hall RA. 1995. Glycosylation and antigenic variation among Kunjin virus isolates. Virology 206:49–56. 10.1016/s0042-6822(95)80018-2.

123. Frost MJ, Zhang J, Edmonds JH, Prow NA, Gu X, Davis R, Hornitzky C, Arzey KE, Finlaison D, Hick P, Read A, Hobson-Peters J, May FJ, Doggett SL, Haniotis J, Russell RC, Hall RA, Khromykh AA, Kirkland PD. 2012. Characterization of virulent West Nile virus Kunjin strain, Australia, 2011. Emerg Infect Dis 18:792-800. 10.3201/eid1805.111720.

124. Haddow AD, Schuh AJ, Yasuda CY, Kasper MR, Heang V, Huy R, Guzman H, Tesh RB, Weaver SC. 2012. Genetic characterization of Zika virus strains: geographic expansion of the Asian lineage. PLoS Negl Trop Dis 6:e1477. 10.1371/journal.pntd.0001477.

125. Shirato K, Miyoshi H, Goto A, Ako Y, Ueki T, Kariwa H, Takashima I. 2004. Viral envelope protein glycosylation is a molecular determinant of the neuroinvasiveness of the New York strain of West Nile virus. J Gen Virol 85:3637–3645. 10.1099/vir.0.80247-0.

126. Moudy RM, Zhang B, Shi P-Y, Kramer LD. 2009. West Nile virus envelope protein glycosylation is required for efficient viral transmission by Culex vectors. Virology 387:222–228. 10.1016/j.virol.2009.01.038.

127. Wen D, Li S, Dong F, Zhang Y, Lin Y, Wang J, Zou Z, Zheng A. 2018. N-glycosylation of Viral E Protein Is the Determinant for Vector Midgut Invasion by Flaviviruses. mBio 9:10.1128/mbio.00046-18. 10.1128/mbio.00046-18.

128. Cimica V, Galarza JM, Rashid S, Stedman TT. 2021. Current development of Zika virus vaccines with special emphasis on virus-like particle technology. Expert Review of Vaccines 20:1483–1498. 10.1080/14760584.2021.1945447.

129. Lin M-H, Li D, Tang B, Li L, Suhrbier A, Harrich D. 2022. Defective Interfering Particles with Broad-Acting Antiviral Activity for Dengue, Zika, Yellow Fever, Respiratory Syncytial and SARS-CoV-2 Virus Infection. Microbiology Spectrum 10:e03949-22. 10.1128/spectrum.03949-22.

